# Mfd affects global transcription and the physiology of stressed *Bacillus subtilis* cells

**DOI:** 10.1101/2020.11.27.401687

**Authors:** Holly Anne Martin, Anitha Sundararajan, Tatiana Ermi, Robert Heron, Jason Gonzales, Kaiden Lee, Diana Anguiano-Mendez, Faye Schilkey, Mario Pedraza-Reyes, Eduardo A. Robleto

**Author notes:** Correspondence: Eduardo A. Robleto.

## Abstract

For several decades, Mfd has been studied as the bacterial transcription-coupled repair factor. However, recent observations indicate that this factor influences cell functions beyond DNA repair. Our lab recently described a role for Mfd in disulfide stress that was independent of its function in nucleotide excision repair and base excision repair. Because reports showed that Mfd influenced transcription of single genes, we investigated the global differences in transcription in wild-type and *mfd* mutant growth-limited cells in the presence and absence of diamide. Surprisingly, we found 1,997 genes differentially expressed in Mfd^-^ cells in the absence of diamide. Using gene knockouts, we investigated the effect of genetic interactions between Mfd and the genes in its regulon on the response to disulfide stress. Interestingly, we found that Mfd interactions were complex and identified additive, epistatic, and suppressor effects in the response to disulfide stress. Pathway enrichment analysis of our RNASeq assay indicated that major biological functions, including translation, endospore formation, pyrimidine metabolism, and motility, were affected by the loss of Mfd. Further, our RNASeq findings correlated with phenotypic changes in growth in minimal media, motility, and sensitivity to antibiotics that target the cell envelope, transcription, and DNA replication. Our results suggest that Mfd has profound effects on the modulation of the transcriptome and on bacterial physiology, particularly in cells experiencing nutritional and oxidative stress.

## 1 Introduction

Short for <u>M</u>utation <u>f</u>requency <u>d</u>ecline, Mfd is the transcription-coupling repair factor, which coalesces a stalled RNAP with the Nucleotide Excision Repair (NER) pathway to preferentially repair lesions in the template strand of actively transcribed genes before lesions in the coding strands or in non-actively transcribed genes. However, as early as the 1990s, evidence suggested that Mfd influenced phenotypes unrelated to transcription-coupled repair. Reports showed that Mfd affected carbon catabolite repression of operons (Zalieckas et al., 1998b). In addition, *in vitro* studies showed that Mfd can facilitate repression transcription by roadblock clearance at genes regulated by the global transcription regulator CodY (Belitsky and Sonenshein, 2011). Our lab demonstrated a role for Mfd in the expression of amino acid biosynthesis genes and protection from oxidative stress (Pybus et al., 2010;Martin et al., 2011;Martin et al., 2019). Of note, a recent report proposed that Mfd functions at hard-to-transcribe genes and affected gene expression and survival associated with toxin-antitoxin gene modules in *B. subtilis* (Ragheb and Merrikh, 2020).

In bacterial species other than *B. subtilis,* observations suggest that Mfd influences traits other than DNA repair. While in some organisms Mfd has the potential to increase the frequency of mutations conferring resistance to antibiotics (Han et al., 2008;Merrikh and Kohli, 2020), in other bacteria such as *H pylori*, it can increase antibiotic sensitivity (Lee et al., 2009). In *Staphylococcus aureus*, inactivation of *mfd* resulted in decreased biofilm formation (Tu Quoc et al., 2007). Interestingly, *E. coli* cells can perform transcription-coupled repair through an Mfd-independent mechanism and the activity of UvrD (Epshtein et al., 2014). This Mfd-independent transcription-coupled repair has been postulated to occur in *B. subtilis* (Moreno-Del Alamo et al., 2020). It has also been shown that genetic lesions of oxidative nature modulate growth and stationary-phase associated Mfd-dependent mutagenic events in this Gram-positive microorganism (Leyva-Sánchez et al., 2020). Moreover, recent single-molecule resolution experiments showed that Mfd can translocate on undamaged DNA independently of its interactions with RNAP (Ho et al., 2018). Given the previous biochemical observations, and the different phenotypes in different bacterial species associated with Mfd, we hypothesized that Mfd, in addition to mediating transcription-coupled repair, modulates the cell transcriptome. More specifically, we tested whether Mfd affects the global transcription profile in stationary-phase *B. subtilis* cells and in conditions of diamide exposure. Diamide is an oxidizing agent that transforms protein thiol groups into disulfide bonds and subject cells to protein oxidation stress (Pother et al., 2009). Cells experiencing disulfide stress activate the oxidative and electrophile stress stimulon to repair and process cellular damage (Antelmann and Helmann, 2011). The activation of this stimulon is complemented and overlaps with regulons controlled by SigB (general stress response), PerR (peroxide stress), OhrR (organic peroxide stress), AdhR (formaldehyde stress), and YodB (disulfide stress) to produce factors that detoxify metabolic intermediates and reactive oxygen species, prevent protein damage, and protect DNA from accumulating lesions (Antelmann and Helmann, 2011).

Using RNA extracted from cultures obtained from stationary-phase *B. subtilis* cultures exposed to either 0 or 1 mM of the protein oxidant diamide, we found that nearly half of the genes in the transcriptome and several biological functions were expressed differentially in the absence of Mfd. Pathway enrichment analysis indicated that the absence of Mfd dysregulates expression of genes affecting biological processes that include translation, endospore formation, and flagellar motility. This dysregulation associated with phenotypic changes in growth in a defined medium, swarming motility, and sensitivity to antibiotics. In addition to identifying pathways affected by Mfd expression, we were also interested in investigating potential gene targets that work in concert with Mfd to confer protection against disulfide stress. Genetic interaction experiments showed that the effect of Mfd on the cellular response to diamide was complex. For example, disruption of *sodA*, which encodes superoxide dismutase, abrogated the ability of cells to survive exposure to diamide, but overexpression of *mfd* in the SodA^-^ background restored it. Our genetic interaction assays uncovered, additive, epistatic, and suppressor effects on the response to disulfide stress.

In conclusion, the results presented here expand the roles of Mfd in the cell beyond transcription-coupled repair and suggest that this factor is a global modulator of transcription with profound effects on bacterial physiology and adaptation to stress.

## 2 Materials and Methods

### Bacterial Strains and Growth Conditions

The parental strain, YB955, is a prophage- “cured” *B. subtilis* strain 168 derivative that contains the point mutations *metB5*, *hisC952*, and *leuC427*. *B. subtilis* strains employed in this study (Table 1) were routinely isolated on tryptic blood agar base (TBAB) (Acumedia Manufacturers, Inc., Lansing, MI, USA), and liquid cultures were grown in Penassay broth (PAB) (antibiotic medium 3, Difco Laboratories, Sparks, MD, USA) supplemented with 1X Ho-Le trace elements (1994). When required, tetracycline (Tet; 10 μg∙mL^−1^), spectinomycin (Sp; 100 μg∙mL^−1^), ampicillin (Amp; 100 μg∙mL^−1^), chloramphenicol (Cm; 5 μg∙mL^−1^), erythromycin (Em; 1 μg∙mL^−1^) or isopropyl-β-D-thiogalactopyranoside (IPTG; 1 mM) were added to media.

**Table 1.**
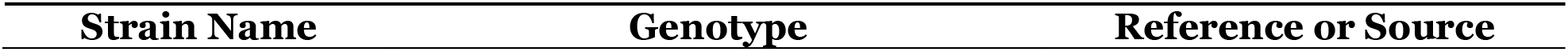
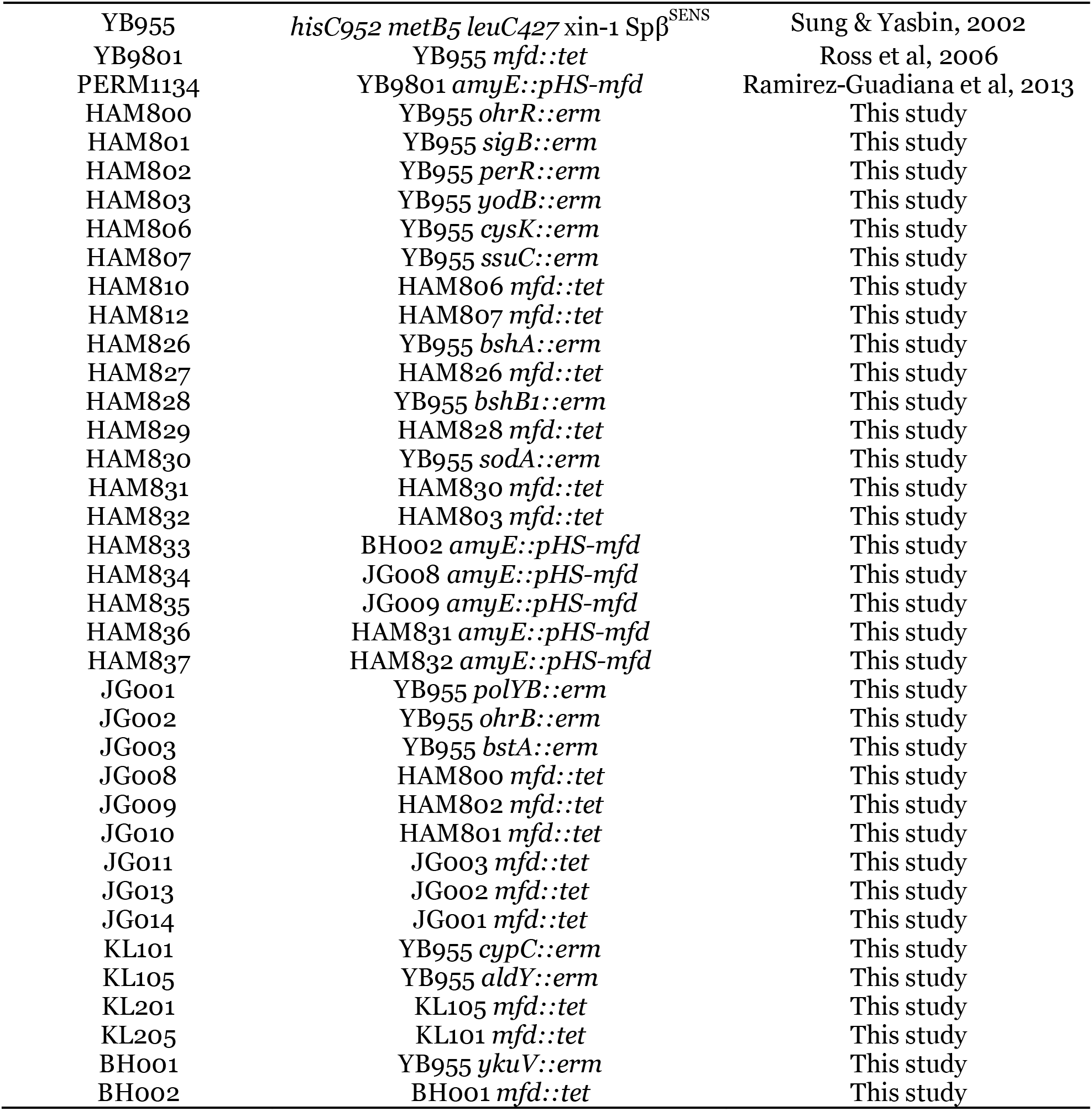
Strains and plasmids used in this study.

### Construction of Mutant Strains

To construct single mutant strains, genomic DNA was isolated from the corresponding BKE (Bacillus Knockout Erythromycin collection - (Koo et al., 2017)) strains using the Wizard® Genomic DNA Purification Kit (Promega, Madison, WI). Of note, the BKE gene deletion constructs are designed to minimize functional interference on the flanking open reading frames. Isolated genomic DNA was then transformed into YB955 using the competence procedures for *Bacillus* described previously (Yasbin et al., 1975). Briefly, YB955 was grown to T_90_, ninety minutes after the cessation of growth (stationary phase), in GM1 broth (0.5% dextrose, 0.1% yeast extract, 0.2% casein hydrolysate, essential amino acids 50 μg/mL, 1X Spizizen salt solution and then diluted 10-fold into GM2 broth (GM1 broth plus 50 μM CaCl2, 250 μM MgCl2). After one hour of incubation at 37°C with aeration, genomic DNA (100 ng) was added. Cells were plated on TBAB containing 5 μg/mL erythromycin to select for the BKE allele. Transformants were confirmed by PCR.

To construct double mutant strains, genomic DNA from YB9801 (Mfd^-^) was isolated and transformed into *B. subtilis* strains with single mutations as described above (Table 1). Cells were plated on TBAB containing 10 μg/mL tetracycline to select for the *mfd^-^* allele and 5 μg/mL erythromycin for maintenance of the BKE allele. Transformants were confirmed by PCR with specific oligonucleotide primers.

To construct the *mfd*-restored strains, BKE or PERM1134 DNA was isolated and transformed as described above. Of note, in this construct, the *mfd* gene is expressed from an IPTG-dependent promoter, and previous experiments showed that IPTG amendment results in restoration of Mfd functions to levels above those observed in the parent strain (Martin et al., 2019). Cells were plated on TBAB containing 100 μg/mL spectinomycin, 10 μg/mL tetracycline, and 5 μg/mL of erythromycin. Transformants were confirmed by PCR.

### RNA sequencing and Differential gene expression analysis

Briefly, a single colony was used to start a two-mL PAB overnight culture. The next morning 0.5 mL was used to start a 15 mL PAB culture. Cultures were grown in flasks containing PAB and Ho-Le trace elements with aeration (250 rpm) at 37 °C until 90 min after the cessation of exponential growth (designated T_90_ (90 min after the time point in the culture when the slopes of the logarithmic and stationary phases of growth intercepted). Growth was monitored with a spectrophotometer measuring the optical density at 600 nm (OD_600_). At T_90_, cultures were divided, and half were treated with 1 mM diamide, and incubated for another two hours.

Total RNA from three biological replicates was harvested from cells differing in Mfd proficiency and treated or untreated with diamide, using the MP FastRNA Pro Blue Kit, and treated with DNase to remove residual DNA (Waltham, MA). Ribosomal RNA was removed by Ribo-Zero Magnetic Kit for Gram-Positive Bacteria, and the remaining RNA was then fragmented. The RNA samples were reverse transcribed into cDNA and sequenced. High quality sequence reads were generated using a HiSeq platform (2×150bp read length) and aligned using HISAT2 (v 2.1.0) short read aligner to the latest version of reference in the Pubmed database (GCA_000009045.1). All sequence data were deposited on SRA under the bio project ID PRJNA673980. Read counts were generated using featureCounts (v1.6.2). Gene expression was quantified as the total number of reads uniquely aligning to the reference, binned by annotated gene coordinate. Differential gene expression and related quality control analyses was determined using the Bioconductor package DESEQ2. Normalization of raw read counts was performed by a scaling method implemented within DESEQ2 package, which accounts for differences in sequencing depth and composition. Differential expression of pairwise comparisons (of the different conditions) was assessed using the negative binomial test with a Benjamini–Hochberg false discovery rate (FDR) adjustment applied for multiple testing corrections.

### Growth assays in complex and defined media

A single colony was used to start a two-mL PAB overnight culture. To start cultures for the growth curve, the OD_600_ for each overnight culture was measured. Cells were diluted to an OD_600_ of 0.4 for each strain and replicate. Then in a 96-well flat-bottom plate, 200 μL of PAB, 10 μL of the diluted overnight cultures, and either 0 mM or 0.5 mM diamide were mixed. Complemented strains were supplemented with 1 mM IPTG to induce expression of *mfd*. The growth curve was incubated at 37° C with shaking on the Synergy HTX plate reader. Readings were taken every five minutes for 16 hours. Each strain was replicated at least nine times.

### Motility assays

A single colony was used to start a two-mL PAB overnight culture. The next morning, the OD_600_ for each overnight culture was measured. Cells were diluted to an OD_600_ of 0.1 for each strain and replicate. 10 μL of the cell dilution was spotted in the middle of TBAB plates containing (0.7% agar) and incubated at 37° C lid-side up, as previously described (Patrick and Kearns, 2009). Plates were examined and photos were taken daily. At least three biological replicates were completed.

### Minimal inhibitory concentration assays

BioMerieux antibiotic strips containing a gradient of concentrations were used to test the effects of Mfd on sensitivity to linezolid, ampicillin, rifampicin, trimethoprim, and daptomycin. Protocols for preparation of cultures and media were followed according to manufacturer’s instructions.

### Statistical analysis

ANOVA was used to test for differences between means. When ANOVA indicated statistical significance between treatments, we used the Least Significant Difference (LSD) method at P< 0.05. ANOVA (complete randomized design) and LSD analyses were conducted using the IBM SPSS 27 software and the statistical package in the GraphPad Prism 8 graphing software.

### Pathway Analysis by Gene Enrichment

Gene enrichment or pathway analyses was performed using the ClueGO plug-in module of the Cytoscape software program, which annotates a list of genes to biological functions (gene ontologies) in a hierarchical way against an annotated genome (Bindea et al., 2009). Lists of genes down or up regulated in the absence of Mfd and in conditions of diamide exposure were used as input into Cytoscape and analyzed for overrepresented gene ontologies. Also, Kappa statistics, within the ClueGO plug-in, were calculated to link gene networks. For example, the list of genes that were downregulated mapped to 13 gene ontology terms and were grouped into 3 groups based on Kappa scores (translation, cell differentiation, and protein folding (Supplementary material). These grouping results were obtained using the ClueGO plug-in of Cytoscape with the following settings: biological functions for gene ontologies, medium network specificity, GO tree interval with level 3 as minimum and level 8 as maximum, enrichment (right -sided hypergeometric test) with a pV value of 0.05 or less with the Bonferroni correction, and a GO term/pathway network connectivity (Kappa Score threshold) of 0.4.

## 3 Results

### Mfd modulates global transcription in stationary-phase and during disulfide stress in *B. subtilis* cells

We conducted transcriptomic analysis assays in stationary-phase cells untreated or treated with 1 mM diamide of the parental (YB955) and Mfd^-^ (YB9801) strains. We used three independent cultures for each condition, which totaled 12 independent observations. The overall results showed that sufficient depth of coverage was established with reads mapping to genes uniquely (mapping to a single location), that expression patterns between independent cultures clustered according to experimental conditions. The results also showed that almost half of the genome was differentially regulated by Mfd and as affected by diamide exposure. This result was striking (Fig. 1A–C). Interestingly, the pairwise comparison between the parent and the Mfd mutant in the absence of diamide exposure showed that mRNA levels of a significant number of genes were dysregulated in the absence of Mfd (Fig. 1C). These results suggested that Mfd has a major impact on the transcription profile of the cell in stationary-phase conditions. Pairwise comparisons for changes in gene expression between the parent strain and the Mfd^-^ mutant in untreated and diamide-treated cells are presented in Table S1.

**Figure 1.**
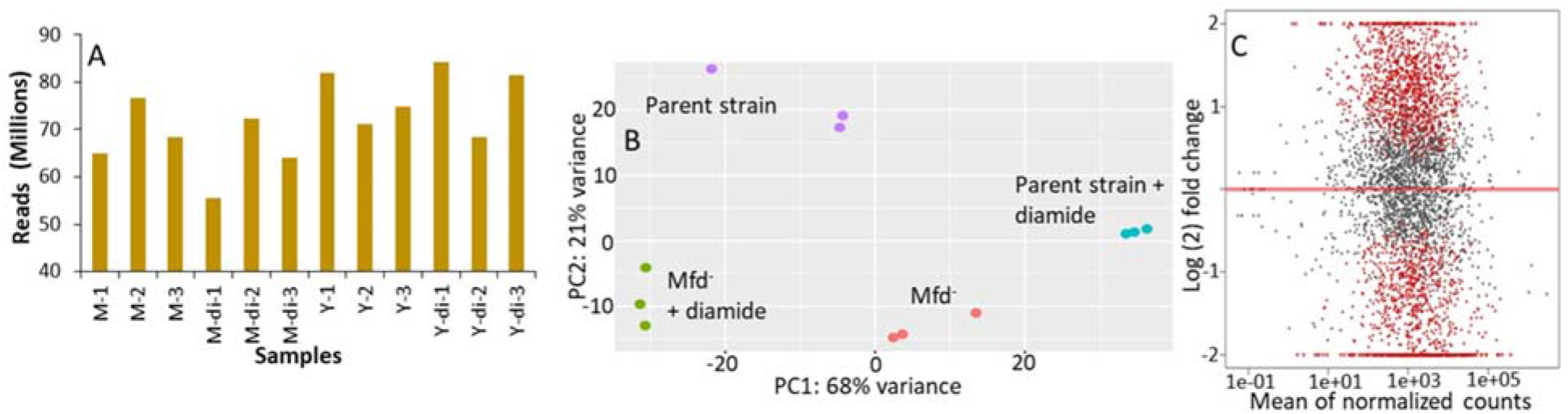
Mfd alters the transcriptome in stationary-phase cells in the presence and absence of diamide. A) Mapped sequencing reads generated by independent RNA samples from M: *mfd* mutant, Y9801; Y: YB955, the parental strain; and di: exposure to diamide. B) Principal component analysis of gene expression results from independent samples (n=3 / strain and treatment combination). C) Gene expression pairwise comparison between the parent, YB955, (reference) and the *mfd* mutant, YB9801, in untreated cells. Each dot represents fold change value in expression of a single gene. Red dots indicate significant change in expression.

In the untreated condition, 1066 genes were downregulated in the absence of Mfd. The values in fold change expression (Log(_2_) in these genes were from −3.55 to −0.39. Pathway-enrichment analysis showed that genes in thirteen biological functions were downregulated in the absence of Mfd. Three major biological pathways were highlighted by Kappa scores: protein folding (Gene Ontology ID 006457), cell differentiation (GO ID 0030154- including spore synthesis and germination), and translation (GO ID 0006412) (Table S2, Fig. S5). On the other hand, expression of 931 genes was upregulated, and fold expression (Log_2_) values ranged from 0.42 to 7.78. Thirty-six biological functions were disproportionately upregulated in the absence of Mfd. These 36 biological functions were coalesced into 10 major groups by Kappa scores and include pentose metabolism (GO ID 0019321), cellular nitrogen and organonitrogen compound biosynthetic processes (GO ID 19001566, GO ID 004271), phosphorus metabolism (GO ID 0006793), toxin metabolism (GO ID 0009404), and transport of carbohydrates and organics (GO ID 0008643 and 0071702), ribonucleoside monophosphate and pyrimidine-containing compound biosynthesis (GO ID 0009156 and GO ID 0072528), and flagellum dependent motility (GO ID 0001539) (Table S3, Fig. S6).

In the diamide treated condition, Mfd deficiency resulted in the down-regulation of 1365 genes. Enrichment pathway analysis showed that genes in 23 gene ontologies were down regulated in the absence of Mfd. These biological functions were grouped by Kappa scores into the ones observed in the untreated condition, proteolysis (GO ID 0006508) and glutamine biosynthesis (GO ID 0009084), isoprenoid metabolism (GO ID 0006720), antibiotic metabolism (GO ID 0016999), carboxylic acid metabolism (GO ID 0019752), and protein metabolism (GO ID 0019538). The fold change in gene expression (Log_2_) ranged from −0.4 to −6.1 (Table S4, Fig. S7). In addition, the Mfd^-^ background displayed up-regulation of 1040 genes, the enrichment analysis showed 24 gene ontologies that included inosine monophosphate (IMP) biosynthesis (GO ID 0006188), cell projection organization (GO ID 0030030), flagellum motility and chemotaxis (GO ID 0001539, GO ID0006935), transmembrane and sodium transport (GO ID 0006814 and 0055085) and cellular nitrogen biosynthesis (GO ID 0044271) (Table S5, Fig. S8. In summary, these results showed that Mfd affects many biological processes in stationary-phase *B. subtilis*.

### Mfd influences sensitivity to antibiotics, growth in defined medium, and swarming motility

Transcriptomic results indicated that loss of Mfd caused dysregulation of global gene expression and prompted us to test for growth and other phenotypes associated with the biological functions that were highlighted by the pathway enrichment analysis (ribonucleoside phosphate biosynthesis, and motility). Mfd affected expression of *rpoB* and *rpoE*, which encode the β and Δ subunits of the RNA polymerase (Boor et al., 1995). The genes *liaF*, *liaR*, and *liaS*, which are activated during oxidative and cell-envelop stress, were differentially expressed by the loss of Mfd (Radeck et al., 2017). Given these observations, we were motivated to investigate the ability to grow on a defined medium (Spizizen medium) and measure minimal inhibitory concentrations (MIC) for ampicillin, linezolid (controls), rifampicin (transcription), daptomycin (cell membrane), and trimethoprim (thymidine synthesis) in cultures of the parent, Mfd^-^, and Mfd^-^ carrying an *mfd*-overexpressing construct in the *amyE* chromosomal locus.

We measured growth in defined medium for 16 hours and showed similar values for doubling time, cell density, and growth lag between the parent and Mfd^-^ cells (Fig. 2A). However, the *mfd*-overexpressing strain showed marked differences in growth lag and cell density compared to the parent or Mfd^-^ cells (Fig. 2A). To better quantify overall growth dynamics, which integrate doubling time and cell density, we used the area under the curve, as calculated by the R-based program GrowthCurver (Sprouffske and Wagner, 2016), to measure the effect of Mfd on growth. The area under the curve (AUC) mean values indicated that the *mfd*-overexpressing cells displayed a significant increase in growth compared to the parent, and the parent’s AUC was significantly higher than the Mfd^-^ strain (Fig. 2B).

**Figure 2.**
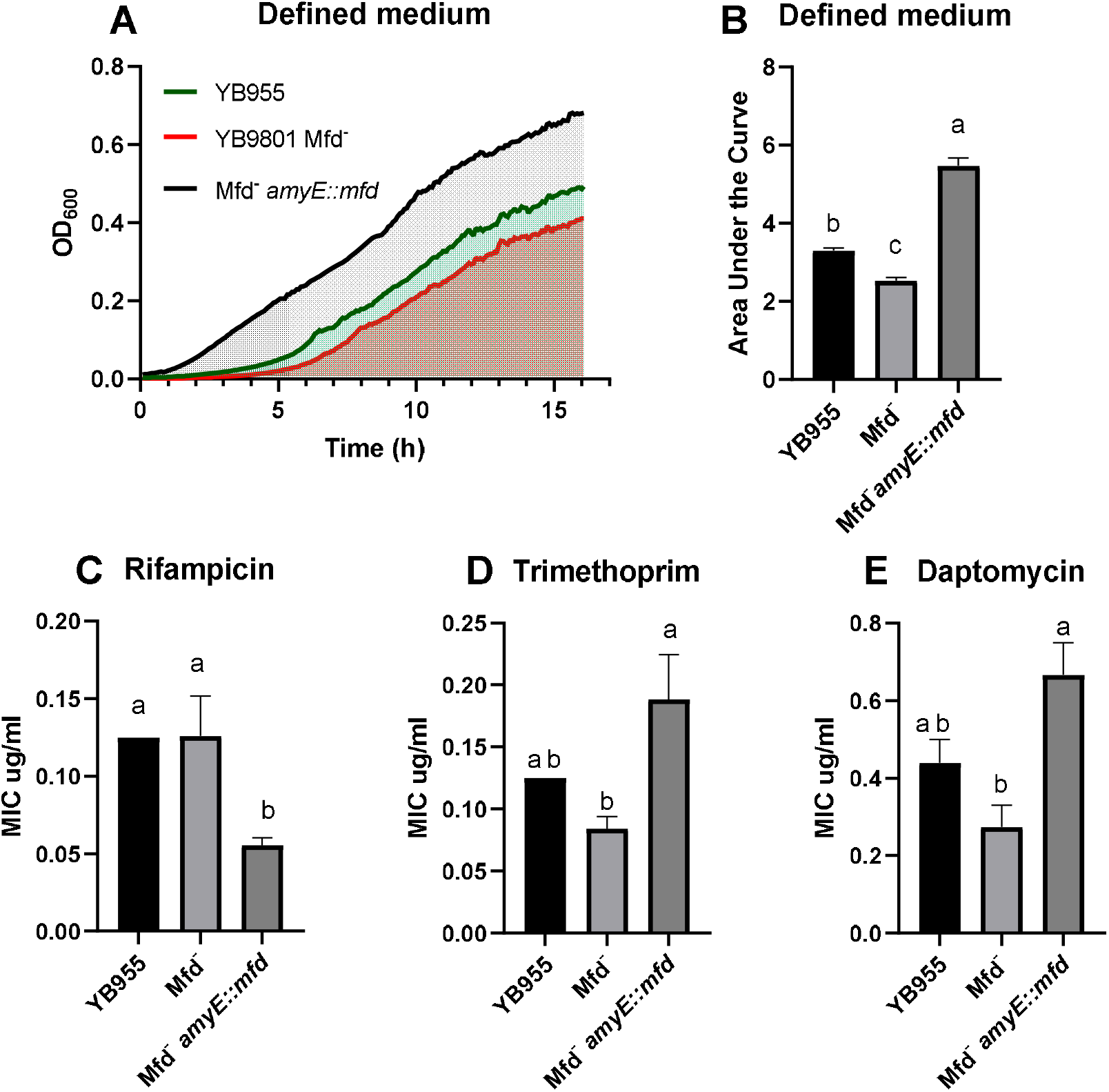
Effects of Mfd on growth in defined medium and minimal inhibitory concentrations MIC to antibiotics. (A) Average OD_600_ readings of strains differing in Mfd measured by a plate reader (n=9). (B) Average area under the curve as calculated by the R package, GrowthCurver, of growth curves from A. (C-E) Average MIC of strains differing in Mfd determined by ETEST strips (n=3). Lower case letters distinguish significant differences between means. “a”, “b”, and “c” are significantly different mean groups.

All the strains were equally sensitive to ampicillin (lowest concentration - 0.016 μg/ml) and resistant to linezolid (highest concentration – 256 μg/ml). Interestingly, the loss of Mfd resulted in lower, but not significant, MIC values than the parent for trimethoprim and daptomycin (Fig. 2D–E). However, overexpression of *mfd* produced significant increases in MIC values when compared to the Mfd^-^ strain (~two-fold and ~three-fold for trimethoprim and daptomycin, respectively). On the other hand, Mfd^-^ cells did not show significant differences to rifampicin when compared to the parent, but the cultures that overexpressed *mfd* were more sensitive than the parent (Fig. 2C).

Cells lacking Mfd showed upregulation of 15 genes (*flgC, flgK, flgL, flhO, flhP, fliE, fliI, fliJ, fliK, fliL, fliM, fliY, hag, motA, and motB -* Table S1) controlling flagellum motility, which prompted us to test if there was a phenotype difference in the swarming ability of *B. subtilis* differing in Mfd. Briefly, overnight cultures were diluted and 10 μL of the cell dilution was spotted in the middle of a TBAB plate, and incubated at 37° C. Each day the colonies were photographed. Cells lacking Mfd displayed increased swarming after two days, and there was further increase after four days. The parent cells and those that overexpressed *mfd* showed reduced swarming (Fig. 3, Fig. S1). Altogether, these results suggest that Mfd affects fundamental aspects of bacterial physiology.

**Figure 3.**
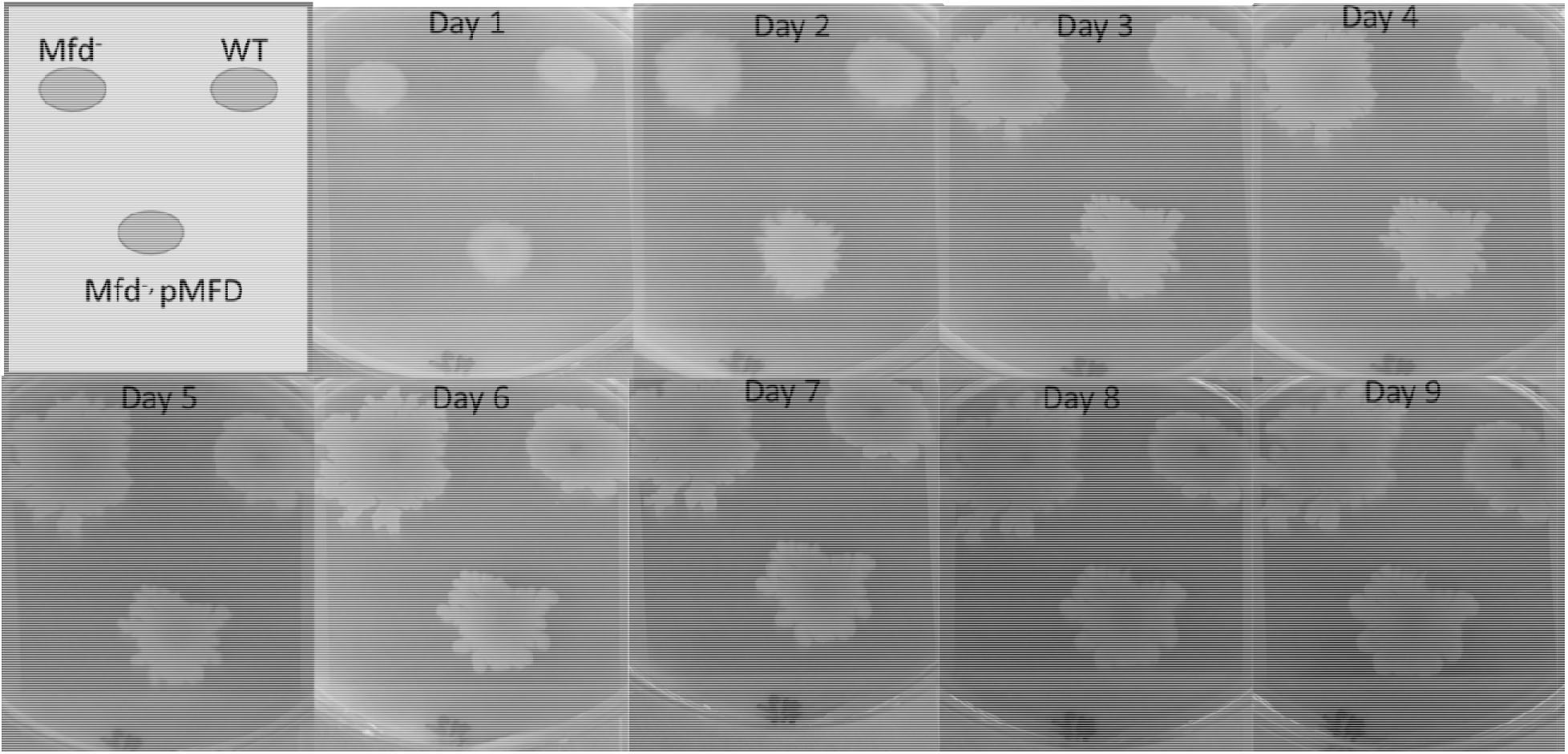
Mfd represses swarming motility. Representative images of swarming of strains differing in Mfd over eight days. Images of additional replicates are available in the supplementary materials.

### Mfd modulates the response to disulfide stress via complex genetic interactions in *B. subtilis*

In addition to the experiments measuring growth in defined medium, sensitivity to antibiotics, and swarming motility, we examined the effect of Mfd in stressed cells and in the context of protein oxidation. Our previous report showed that Mfd protected cells against disulfide stress, but not much is known about how this factor operates to provide cells such protection. Growth in complex broth in the absence and presence of diamide was followed by OD_600_ measurements every five minutes for 16 hours. Interestingly, the growth dynamics of the parent and Mfd^-^ strains were indistinguishable in the untreated cells, and both genetic backgrounds showed a similar growth lag in the presence of diamide (Fig. 4A). However, the Mfd^-^ cells displayed an increase in doubling time during mid exponential growth and a decrease in cell density compared to the parent strain in the presence of diamide. Cells that overexpressed *mfd* displayed the same growth rate but an increase in cell density compared to the parent strain in untreated conditions (Fig. 4A). Strikingly, the *mfd*-overexpressing cells showed a shorter growth lag and a higher cell density than the parent strain, but both strains showed a similar doubling time in the treated cells (Fig. 4A). The AUC values showed that the growth responses in untreated cells were significantly different (P< 0.05) and that the *mfd*-overexpressing cells responded better than the parent and Mfd^-^ counterparts (Fig. 4B). Overexpressing *mfd* increased tolerance to diamide levels significantly better than in the parent and Mfd^-^ counterparts. In fact, the response in the *mfd*-overexpressing strain was like the one observed in the untreated parent strain. The Mfd^-^ cells displayed the lowest tolerance to diamide amongst the tested strains (Fig. 4B). The results observed in untreated and diamide-treated cells suggest that Mfd controls the physiology of cells experiencing nutrient-limiting conditions and the response to disulfide stress.

**Figure 4.**
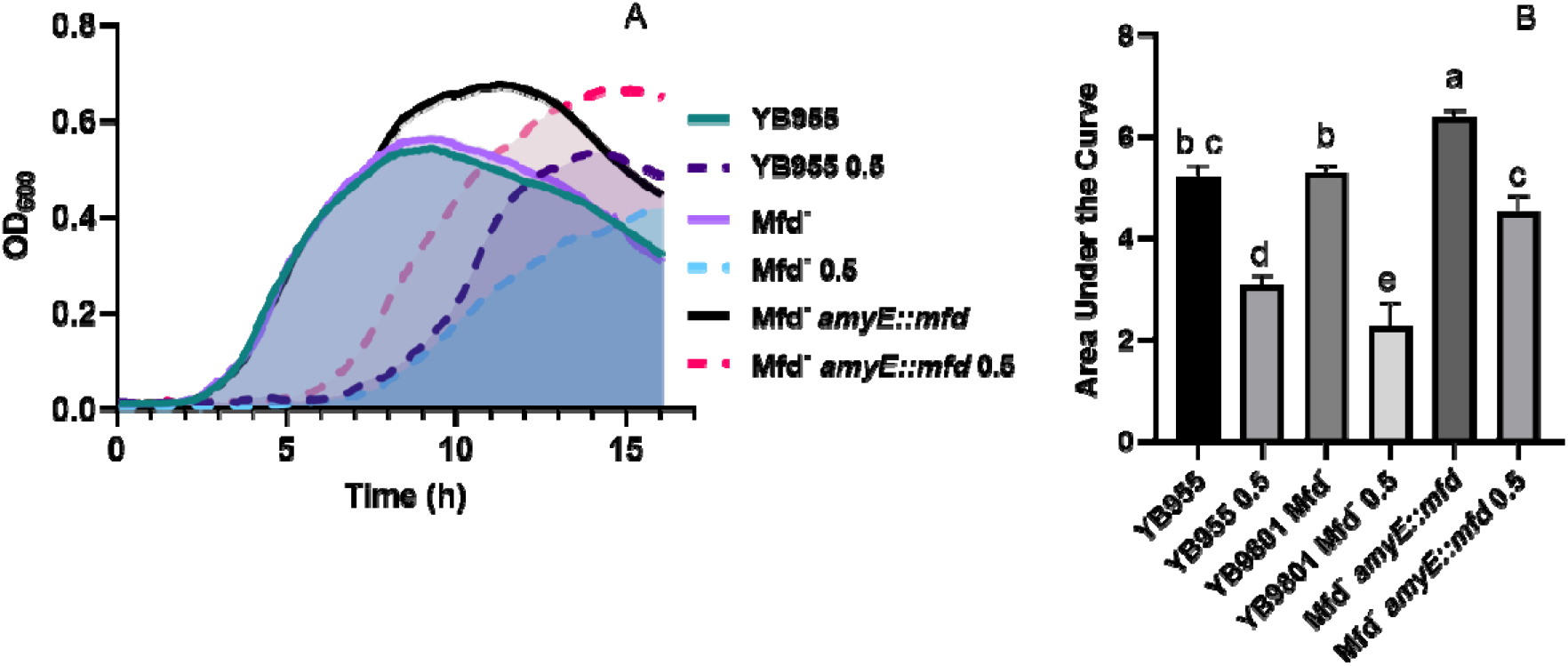
Effects of Mfd on growth in a complex medium and in the presence of diamide. (A) Average OD_600_ readings of strains differing in Mfd measured by a plate reader (n=9). (B) Average area under the curve as calculated by the R package, GrowthCurver, of growth curves from A. Lower case letters distinguish significant differences between means. “a”, “b”, “c” and onward are significantly different mean groups.

To better understand the role of Mfd in the response to disulfide stress, we examined the genetic interactions between the Mfd factor and those genes that are either affected in expression by Mfd, are known transcription factors that control gene expression during oxidative damage, or are components of cysteine and sulfur metabolism, which produce compounds important for disulfide and electrophile stress tolerance (Hochgräfe et al., 2007;Gaballa et al., 2010). We selected 15 genes to generate single knockouts in our parental strain, YB955 (Table 1 and Table S6). Then, we produced double knockouts of Mfd and each of the 15 selected genes by transforming genomic DNA from our Mfd mutant strain, YB9801, into the single knockout. Also, for 5 of the 15 genes under study, we generated strains in which the double knockout was transformed with genomic DNA from our *mfd*-restored strain, PERM1134. This process generated three types of strains: i) single gene knockouts, ii) double knockout combinations of *mfd* and each of the 15 genes, and iii) double knockouts with an *mfd* overexpressing construct ectopically placed in the chromosome.

We conducted growth assays in the presence and absence of 0.5 mM of diamide and used the area under the curve to measure the growth response and tolerance to diamide in all the tested strains. Strains with single mutations in *bshA, bshB1*, and *ohrB* showed a similar growth response and tolerance to diamide to the parent (Fig. 5A–C, Fig. S2). These results suggest that the single contributions of these factors to disulfide stress tolerance are negligible (Fig. 5A–C). However, the mean response in tolerance to diamide in cells with mutations in *mfd* and each of these three factors were statistically the same as the one observed in the parent. These results suggest that mutations in *bshB1*, *bshA*, and *ohrB* suppress the effect of *mfd* on disulfide stress.

**Figure 5.**
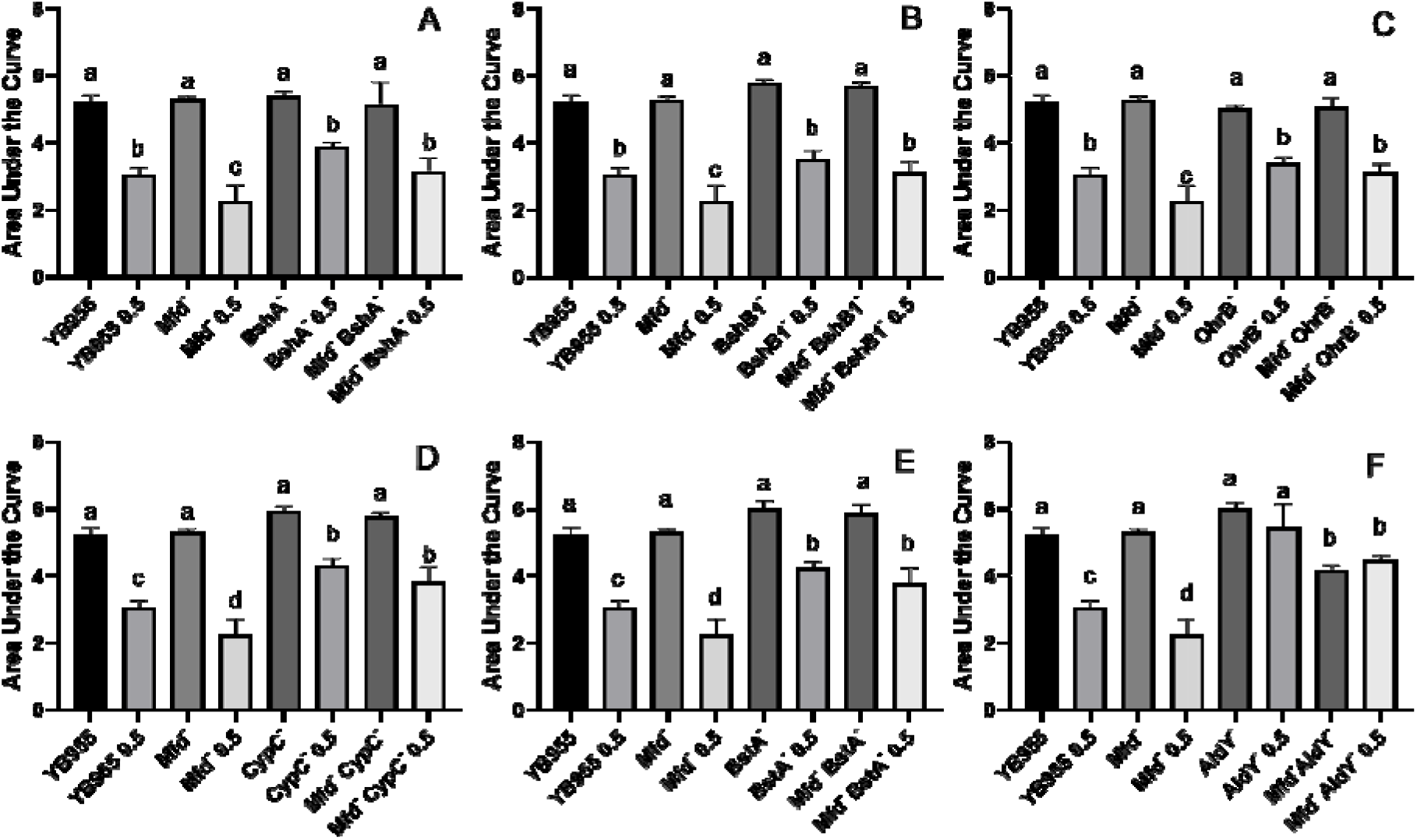
Growth dynamics of single- and double-gene-deletion strains in the presence and absence of diamide show gene suppressors and recessive interactions. Average area under the curve as calculated by the R package, GrowthCurver, of growth curves from Fig S#. Lower case letters distinguish significant differences between means. “a”, “b”, “c” and onward are significantly different mean groups.

Mutations in *cypC*, *bstA*, and *aldY* did not affect growth, but exerted increased tolerance to diamide when compared to the parent or Mfd mutant (Fig. 5D–F, Fig. S2). Interestingly, the levels of increased tolerance to diamide in the CypC^-^ and BstA^-^ backgrounds were unaffected by mutations in *mfd*. These results suggest that the loss of these factors alters the cell physiology to better withstand disulfide stress and that the genes encoding each of these factors exert a recessive epistatic effect on *mfd*. Of note, the combined mutations in *mfd* and *aldY* showed a decreased growth compared to the parent, which suggests that both functions are required for optimal growth, but their absence produces an overall response that desensitizes cells to disulfide stress (Fig. 5F).

We conducted experiments that included overexpression of the Mfd factor in strains with mutations in transcription factors that affect the response to either oxidative or disulfide stress, as well as in strains with mutations in genes coding for a thiol-oxidoreductase and a superoxide dismutase (Inaoka et al., 1998;Fuangthong et al., 2001;Zhang et al., 2006). Mutating *ohrR* did not affect growth, but decreased tolerance to diamide treatment to the levels observed in the Mfd^-^ background (Fig. 6A, Fig. S3A). The results shown by the double inactivation of *ohrR* and *mfd* suggest that these genes contribute to diamide additively and, therefore, are part of different cellular pathways (Fig. 6A). Restoring *mfd* with an overexpressing construct in the OhrR^-^ Mfd^-^ mutant resulted in increased tolerance to diamide levels higher than the ones observed in the parent (Fig. 6A). Deficiencies in YodB, a repressor that controls expression of genes that respond to disulfide stress (Chi et al., 2010), did not affect growth, but displayed decreased tolerance to diamide levels comparable to the Mfd^-^ strain. However, the combined effects of deficiencies in Mfd and YodB resulted in increased growth and tolerance to diamide, and overexpressing *mfd* in this background sensitized cells to it (Fig. 6B, Fig S3B). Inactivation of PerR, the repressor that controls the response to peroxide (Bsat et al., 1998), increased growth but showed no differences in tolerance to protein oxidation compared to the parent strain. The response in the Mfd^-^ PerR^-^ strain suggests that a component of the increase in growth seen in the Per^-^ cells is dependent on Mfd; restoring this factor restored growth to the levels seen in the Per^-^ cells but did not change the response to diamide compared to the double mutant (Fig. 6C, Fig. S3).

**Figure 6.**
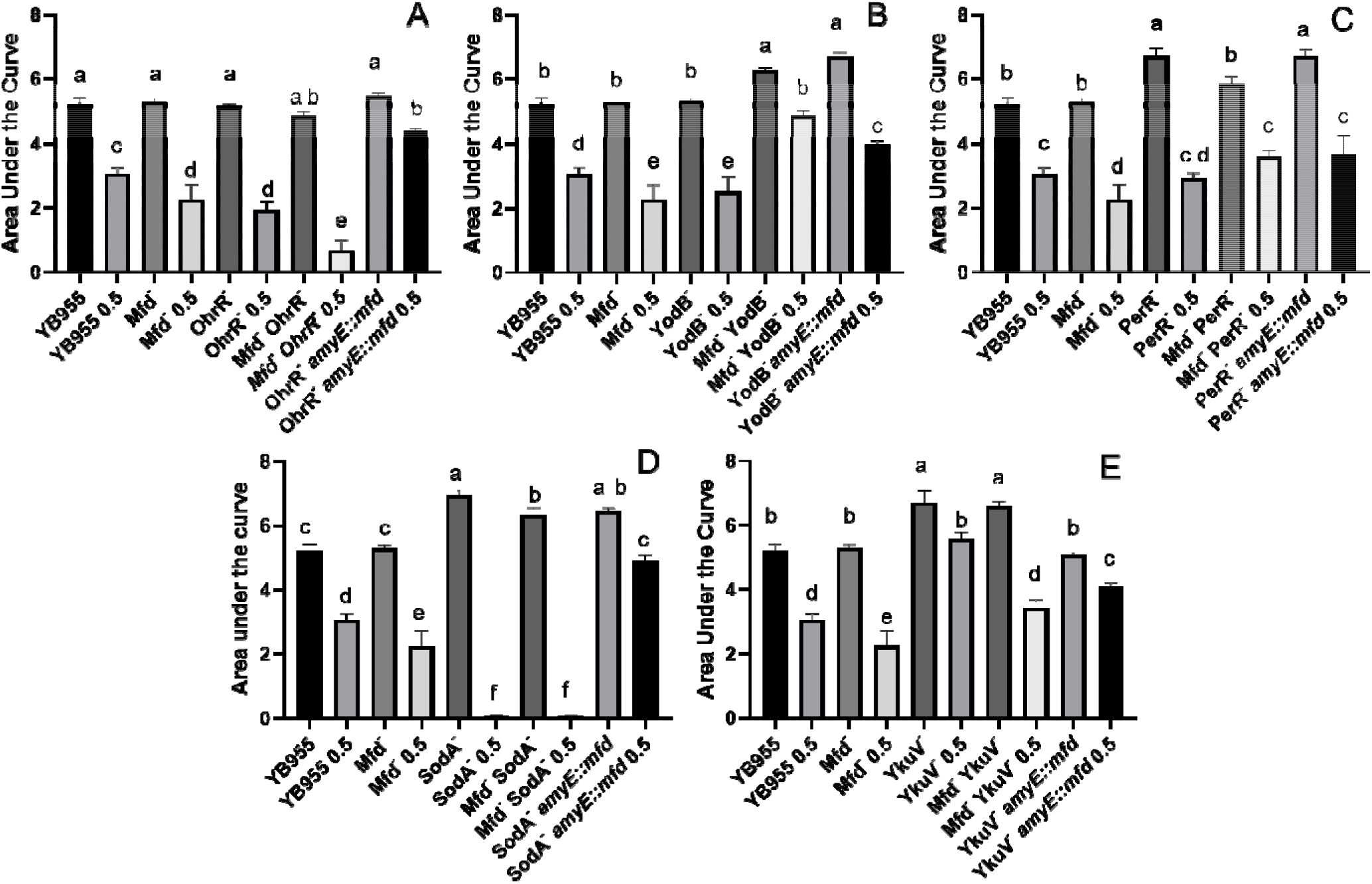
Overexpression of *mfd* can rescue or sensitize cells to diamide. Growth dynamics of single- and double-gene-deletion and overexpressing-*mfd* strains in the presence and absence of diamide. Average area under the curve as calculated by the R package, GrowthCurver, of growth curves from Fig S3. Lower case letters distinguish significant differences between means. “a”, “b”, “c” and onward are significantly different mean groups.

Genetically inactivating SodA increased growth but abrogated tolerance to protein oxidation, and a similar response was observed when this mutation combined with Mfd deficiencies in treated and untreated cells (Fig. 6D, Fig, S3). This result suggests that the *sodA* gene has a recessive epistatic effect onto *mfd*. However, tolerance to diamide was restored to levels higher than the parent in the SodA *-* Mfd^-^ strain that overexpressed *mfd*. This indicates that excess Mfd in the cell can remodel physiology to bypass the absence of SodA. Interestingly, cells that lack YkuV, which encodes a thiol oxidoreductase, showed an increase in growth and tolerance to diamide compared to the parent, and the values in diamide tolerance seen in the double mutant (Mfd^-^ YkuV^-^) suggest a suppressor effect onto the *mfd* mutation. Overexpression of *mfd* restored growth to the parent levels and increased tolerance to diamide significantly compared to the double mutant (Fig. 6E, Sig, S3). This result suggests that excess Mfd can function to increase tolerance to diamide independently of the suppressor effect exerted by the loss of YkuV (compare the means of the double mutant and the YkuV^-^ mutant overexpressing *mfd* in the presence of diamide - Fig. 6E).

The experiments examining the pairwise interactions between *mfd* and *sigB* (general stress response (Petersohn et al., 1999)), *ssuC* (aliphatic sulfonate transporter (van der Ploeg et al., 1998)), *polYB* (translesion synthesis polymerase (Duigou et al., 2004)), and *cysK* (cysteine synthase (van der Ploeg et al., 2001)) showed that single mutations in the latter four genes resulted in an increase in growth and tolerance to protein oxidation (Fig. S4). The double mutants Mfd^-^ SigB^-^, Mfd^-^ PolY^-^, and Mfd^-^ SsuC^-^ showed a similar response to the ones observed in the Mfd^+^ background, which suggests that strains with deficiencies in either SigB, PolYB, or SsuC respond to disulfide stress independently of Mfd. In contrast, the double mutant Mfd^-^ CysK^-^ showed a decreased growth and tolerance to diamide compared to the CysK^-^ cells but expressed similar values in treated and untreated conditions (Fig. S4). This result suggests that combined deficiencies in CysK and Mfd decrease growth but do not influence overall tolerance to diamide. Altogether, these results indicate that the effects of Mfd on the growth and the response to protein oxidation is complex and dependent on gene interactions, and that this factor can remodel cell physiology to sensitize or increase tolerance to disulfide stress.

## 4 Discussion

Here, we examined the effect of Mfd on global transcription, growth in different media, sensitivity to antibiotics, motility, and the response to protein oxidation. Our experiments were prompted by reports suggesting that Mfd affects biological functions other than DNA repair in *B. subtilis* and other bacterial species (Zalieckas et al., 1998b;Tu Quoc et al., 2007;Han et al., 2008;Lee et al., 2009;Pybus et al., 2010). Specifically, in *B. subtilis*, our previous report showed that Mfd protected cells experiencing disulfide stress (Martin et al., 2019).

Mfd affected expression of almost 2000 genes in stationary-phase cells and several biological functions, even in the absence of disulfide stress. While there are reports that demonstrate that this factor affects transcription of single genes controlled by catabolite repression and branch chain amino acid starvation (Zalieckas et al., 1998b;Belitsky and Sonenshein, 2011), there are no reports on the effects of Mfd on overall cell physiology on stationary-phase bacterial cells. Mfd did not affect the growth dynamics in a complex medium compared to the parental strain (Fig. 4). However, Mfd^-^ cells showed differences in expression of 108 essential genes (83 were down regulated in the absence of Mfd – see table S1) that affect DNA replication, cell division, transcription, and protein synthesis. Given these changes in gene expression, we tested Mfd derivatives for their ability to grow in defined medium and sensitivity to antibiotics that target cell envelope, transcription, and DNA replication. Mfd defects did not result in significant changes in MIC for rifampicin, trimethoprim, and daptomycin, but its overexpression did. While Mfd^-^ cells displayed the same MIC values for rifampicin compared to the parent strain, the *mfd*-overexpressing strain was significantly different from the Mfd^-^ mutant and showed significantly lower MIC values than the parent strain. The MICs for trimethoprim and daptomycin were significantly higher in the *mfd*-overexpressing cells than the Mfd^-^ cells. The results in growth in defined medium and antibiotic sensitivity support the idea that the Mfd^-^dependent changes in gene transcription alter cell physiology.

It is possible to speculate that the effect of Mfd on rifampicin MIC is likely due to the changes in expression in *rpoB* and *rpoE* in the Mfd^-^ cells (Table S1). Mfd is active during transcription elongation and at regions distant from the promoter (Haines et al., 2014) and processes paused RNAP into active transcription (Park et al., 2002). Perhaps, excess Mfd alters the equilibrium between the RNAPs engaged in initiation and elongation, and such alteration results in the increased sensitivity observed here. In the absence of Mfd, 65% of the genes for inosine monophosphate metabolism were upregulated. Also, other genes involved in nucleoside metabolism were affected by Mfd. The changes in gene expression of IMP biosynthetic genes are likely to affect nucleotide pools and induce a more sensitive state to trimethoprim; such sensitivity was reduced when *mfd* was overexpressed. Interestingly, activity of the ribonuclotide reductase, encoded by *nrdEF*, influences Mfd^-^dependent mutagenesis (Castro-Cerritos et al., 2017;Castro-Cerritos et al., 2018). The response in daptomycin correlated with low gene expression of the *liaRS* operon, which codes for factors that sense cell-envelope stress. It is worth noting that previous studies have implicated Mfd in the formation of mutations that confer resistance to antibiotics (Han et al., 2008;Lee et al., 2009;Merrikh and Kohli, 2020); however, no reports associate Mfd^-^dependent changes in gene expression to antibiotic sensitivity.

Our assays also showed that the absence of Mfd resulted in increased swarming motility. This result correlated with the RNASeq results that indicated that several genes controlling this process were upregulated in the absence of Mfd. To our knowledge, this is the first report that shows Mfd to be a factor that controls motility. Interestingly, the RNASeq findings indicated that 40% of genes involved in endospore formation, another coordinated cellular behavior, were downregulated and suggest that the loss of Mfd affects sporulation. This assertion agrees with our previous report (Ramirez-Guadiana et al., 2013) and that of Koo et al., which showed that Mfd^-^ deficient cells display a lower sporulation efficiency than Mfd^+^ cells (Koo et al., 2017). The decrease on sporulation efficiency could be the result of the combined deficiencies in gene expression and repair of oxidative damage during endospore formation in Mfd^-^ cells. Mfd^-^ cells with defects in DNA repair systems that target 8-oxo-G showed an exacerbated decrease in sporulation efficiency during oxidant exposure compared to the Mfd^-^ cells (personal communication).

The loss of Mfd significantly decreased growth in defined medium, and overexpressing this factor produced an increase in growth response in defined medium compared to the parent. On the other hand, growth in PAB (complex medium) showed no differences between the parent and the Mfd^-^ cells, but overexpression of Mfd conferred PAB cultures the ability to attain a higher AUC density than the parent. Perhaps, Mfd facilitates gene expression of amino acid biosynthetic genes and a more efficient growth physiology in defined medium; in complex medium, Mfd^-^dependent changes in physiology allow cells to continue doubling during the transition to nutrient scarcity. Interestingly, the experiments that constructed and tested strains containing non-essential gene deletions for their ability to grow in LB in different conditions and amendments showed that Mfd^-^ cells had a slight growth defect in LB at 37° C, as measured by relative fitness (Koo et al., 2017). The results from our growth assays suggest that the biological consequences associated with Mfd express in cells nutritionally stressed or conditions in which amino acid biosynthesis is active.

The effects of Mfd on the overall response to diamide exposure were complex and influenced by different gene interactions. In untreated conditions, several gene ontologies were affected by Mfd, and while there were no consequences in the ability to grow on PAB, it could be argued that the changes in gene expression observed in the untreated condition sensitize cells to protein oxidation. Based on our RNASeq results, cellular functions that were downregulated in the absence of Mfd in treated cells only include, protein degradation, glutamine, isoprenoid and carboxylic acid metabolism, as well as antibiotic metabolism. Contrastingly, genes involved in inosine monophosphate biosynthesis, cell projection and organization, as well as transmembrane and sodium transport were upregulated in the absence of Mfd in treated cells only. So, it can be speculated that such changes in gene expression result in a cellular state that compromise the response to disulfide stress. Future research will examine the specific contributions of those gene ontologies to sensitivity and adaptation to diamide exposure.

To better determine how Mfd affects the response to disulfide stress, we measured the AUC in strains carrying single and pairwise combinations of mutations of *mfd* and genes whose expression was downregulated in the Mfd^-^ background (Fig. 5 and Fig. S4). We also examined genetic interactions between *mfd* and factors involved in oxidative and electrophile stress, as well as sulfur and cysteine biosynthesis. The results indicated different kinds of interactions, and the strains with single mutations in *bshA*, *bshB1*, *ohrB*, *cypC*, *bstA*, *sigB*, and *polYB* showed no significant differences between the parent and Mfd^-^ backgrounds. Therefore, one can argue that the effects on growth in a complex medium and tolerance to diamide caused by the single mutations in these genes (*bshA*, *bshB1*, *ohrB*, *cypC*, *bstA*, *sigB*, and *polYB*) are Mfd^-^independent. Contrastingly, the effects of mutations in *aldY*, *ohrR*, *yodB*, *perR*, *sodA*, *ykuV*, *ssuC*, and *cysK* on growth in PAB (*aldY*, *yodB*, *perR*, *sodA*, and *cysK*) or tolerance to diamide (*aldY*, *ohrR*, *yodB*, *ykuV*, and *cysK*) were partially dependent on Mfd.

The effect of Mfd overexpression on growth and tolerance to diamide was dependent on the genetic background tested (Fig 6). Mfd^-^ PerR^-^ *amyE::mfd* cells grew better in PAB than their PerR^-^ Mfd^-^ counterparts but did not affect the response to diamide exposure. Contrastingly, overexpression of *mfd* combined with mutations in *ohrR* and *sodA* resulted in increased tolerance to disulfide stress when compared to Mfd^-^ OhrR^-^ and Mfd^-^ SodA^-^ cells, respectively. Growth in untreated cells with those genetic backgrounds was not affected. The cells that overexpressed *mfd* in the YodB^-^ background showed a decrease in tolerance to disulfide stress compared to the Mfd^-^ YodB^-^ cells, and overexpression of *mfd* in the YkuV^-^ background resulted in decreased growth and increased tolerance to diamide. While overexpression of Mfd may not be relevant to physiological conditions, we interpreted these results to mean that Mfd can function to modulate the response to diamide by different pathways.

Mfd is a multidomain enzyme that can translocate on DNA independently of its direct interactions with RNAP (Deaconescu et al., 2006;Ho et al., 2018;Le et al., 2018). While most studies on this factor have focused on its transcription-coupled DNA repair functions, recent in vitro studies have shown that Mfd can function as a transcription modulator by reactivating paused RNAP into active transcription or dissociating transcription elongation complexes blocked by lesions or DNA-protein complexes(Selby and Sancar, 1995;Belitsky and Sonenshein, 2011;Le et al., 2018). Also, studies in mutagenesis suggest that Mfd acts directly at highly transcribed regions that accumulate lesions and mediates the formation of mutations (Gomez-Marroquin et al., 2016). Interestingly, other reports showed that Mfd contributes to catabolite repression, and that such repression requires the CcpA protein complexed to the catabolite repression element (*cre*) sequence at promoter distal regions in the *acsA* gene (Zalieckas et al., 1998b). In the absence of Mfd, active transcription at catabolite-repressed genes is blocked by the *cre*-CcpA complex, but not terminated as the paused RNAP is not dissociated from the transcribed DNA (Zalieckas et al., 1998a). Subsequent rounds of transcription may lead to rear-end collisions between active and stalled RNAPs; such collisions could clear the DNA-protein block and lead to resumed transcription. This mode of transcription termination was shown also for genes controlled by the CodY repressor (Belitsky and Sonenshein, 2011). So, it is possible to speculate that Mfd can modulate transcription directly to reduce or increase the levels of complete transcripts of a given gene. If Mfd is required to process RNAP paused at intrinsic sites or pauses caused by non-B DNA structures back into active transcription, then loss of Mfd would result in a decrease of complete gene transcripts. In vitro studies demonstrated that non-B DNA structures can stall transcription (Tornaletti et al., 2008;Pandey et al., 2015). On the other hand, Mfd can terminate transcription directly by dissociating RNAPs that have encountered DNA-repressor protein blocks, and loss of this factor would lead to de-repression of genes and the accumulation of complete transcripts, as suggested by the reports in catabolite and CodY repression (Zalieckas et al., 1998a;Zalieckas et al., 1998b;Belitsky and Sonenshein, 2011). Our RNAseq and phenotypic assays cannot discern between the direct and indirect effects on gene expression caused by the Mfd factor, and based on the number of Mfd molecules per cell (estimated at 30-300 - (Ho et al., 2018)), it is likely that a large fraction of the ~2,000 genes affected by Mfd are caused by indirect effects. However, the experiments testing gene interactions and *mfd* overexpression are informative on which cellular pathways are dependent of Mfd to exert increases and decreases in growth and tolerance to protein oxidation. Given the pleiotropic effects observed in stressed Mfd^-^ cells, Mfd may directly modulate the levels of full-length transcripts of genes that respond to a specific stress or are important for development or cell behavior.

A recent genome association report showed that expression of almost 380 genes were affected by Mfd during exponential growth; genes that were shown to be directly affected by this factor encoded toxin-antitoxin modules (Ragheb and Merrikh, 2020). Further, those studies showed that overexpression of the *txpA*, *bsrH*, genes in a Mfd^-^ cells decreased cell survival compared to the parent strain and led to propose a model in which Mfd directly interacts with RNAP to terminate transcription of regions with structured RNAs (Ragheb and Merrikh, 2020). Interestingly, our RNA-Seq assays, conducted in stationary-phase cells indicated 1997 genes to be affected by Mfd and showed no changes in expression of *txpA* and *bsrH* genes in untreated cells. However, Mfd^-^ cells treated with diamide were downregulated in transcription of those genes (−1.14 Log_(2)_ for *bsrH* and −1.79 for *txpA*), which suggests that Mfd^-^dependent modulation of transcription changes according to growth conditions. The results reported here and those reported in exponentially growing cells provide strong evidence for the concept that Mfd acts as global modulator of transcription and that the biological consequences associated with this factor manifest in stressed cells. Future experiments that measure transcription pauses as affected by Mfd (NETseq) in combination with assays that target DNA regions associated with Mfd (ChIPseq) will further elucidate how Mfd influences cell physiology directly. A recent report used RNETseq to determine NusG-dependent transcription pauses and identified 1600 TTNTTT pause motifs. Of note, expression of NusG was downregulated in the absence of Mfd (−1.25 Log_(2)_ value, Table S1) in this study. Also, our group recently reported roles for GreA (expression of this gene was downregulated; −1..09 Log_(2)_ value, TableS1), a transcription fidelity factor that processes backtracked RNAP, in stress-induced mutagenesis (Leyva-Sanchez et al., 2020). So, it is reasonable to speculate that Mfd can indirectly affect transcription modulation through GreA and NusG.

In summary, this work demonstrates that Mfd has profound effects on the transcriptome and phenotypes of stationary-phase *B. subtilis*, and that such effects do not manifest in cells growing under optimal conditions. The biological consequences associated with Mfd extend beyond deficiencies in transcription-coupled DNA repair and are expressed in cells exposed to nutrient deprivation, cell envelope stress, antibiotics, and protein oxidation. Also, Mfd influences behaviors that include cell differentiation (Ramirez-Guadiana et al., 2013;Koo et al., 2017) (personalcommunication/manuscript in review) and swarming motility. Because Mfd is well conserved in bacteria, it would not be surprising if the pleiotropic effects observed in *B. subtilis* extend to other bacterial species, including those that are pathogenic. Considering the mutagenesis functions of Mfd that confer fitness to stationary-phase cells and resistance to antibiotics, our results suggest that this factor operates at the intersection between gene expression that responds to stress and bacterial evolution.

## 8 Author Contributions

The experiments were planned by HAM, AS, FS, MPR, and EAR. HAM, TE, RH, JG, KL, DAM, and EAR performed the laboratory work. The data was analyzed by HAM, AS, MPR, and EAR. HAM and EAR wrote the manuscript, which was edited by AS, TE, RH, JG, KL, DAM, FS, and MPR. All authors contributed to the article and approved the submitted version.

## 10 Acknowledgements

This research was performed as a collaboration with the New Mexico IDeA Networks for Biomedical Research Excellence (INBREs) supported by an Institutional Development Award (IDeA) from the National Institute of General Medical Sciences of the National Institutes of Health under grant number P20GM103451.

