## Supplementaty materials for "Mfd affects global transcription and the physiology of stressed *Bacillus subtilis* cells": Copy of Diamide Paper Suppl_figs_tables_10_29_20.docx

**Supplementary Materials.**

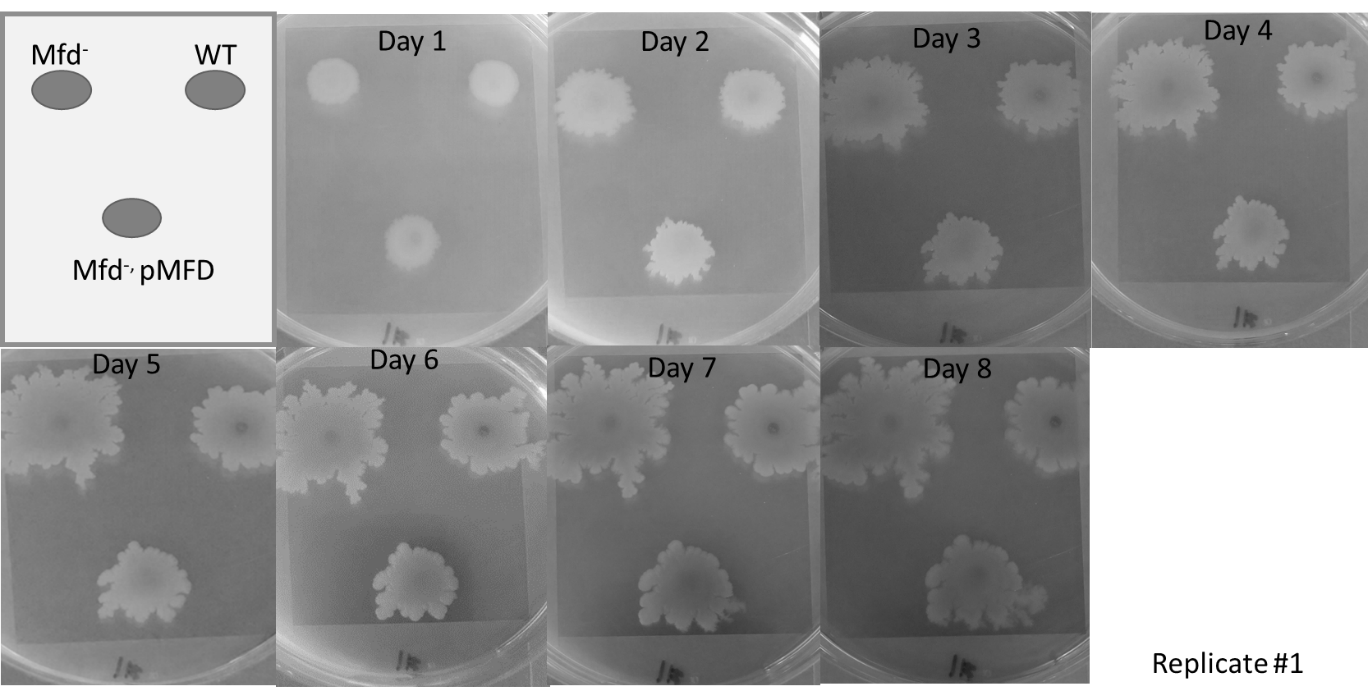

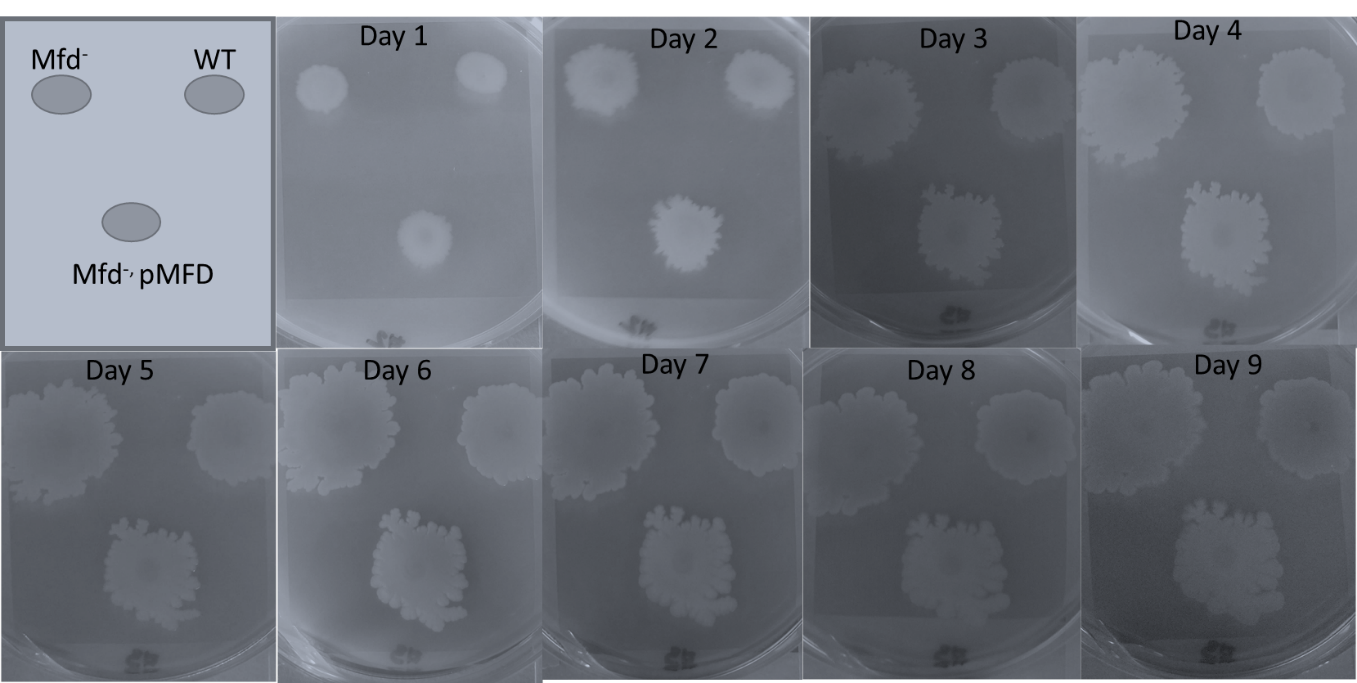

**Figure S1. Additional replicates: Mfd represses swarming motility.** Replicate images of swarming of strains differing in Mfd over eight or nine days.

**Table S6. Description of genes chosen to investigate Mfd genetic interaction and their effects on disulfide stress.**

| **Gene** | **log_2_FoldChange** | **function** | **Regulon** | **Oxidative/Electrophile Stimulon Member** |
| --- | --- | --- | --- | --- |
| *cysK* | -1.335166409 | biosynthesis of cysteine, control of CymR activity | CymR regulon, Spx regulon, SigA regulon, SigM regulon | no |
| *ssuC* | -1.090178022 | sulfonate uptake | CymR regulon, SigA regulon | no |
| *bstA* | -0.974382296 | detoxification; bacillithiol S-transferase | BstA regulon | yes |
| *ohrB* | -1.542418572 | organic peroxide resistance | SigB regulon | yes |
| *polYB (polY2)* | -1.149041837 | UV-targeted mutagenesis | LexA regulon | no |
| *aldY* | -1.356597831 | stress resistance; aldehyde dehydrogenase (NAD) | SigB regulon | yes |
| *cypC* | -1.37096633 | biosynthesis of beta-hydroxy fatty acid for lipopeptides | SigB regulon | yes |
| *ohrR* | n/a | regulation of ohrA expression in response to organic peroxides | SigA regulon, OhrR regulon | yes |
| *sigB* | n/a | general stress response, biocontrol of fungal growth | SigA regulon, SigB regulon, CcpA regulon | no |
| *perR* | n/a | regulation of the response to peroxide | PerR regulon, SigA regulon | yes |
| *yodB* | n/a | regulation of quinone and diamide detoxification | YodB regulon | yes |
| *sodA* | -1.06855698 | detoxification of oxygen radicals | SigB regulon | yes |
| *bshA* | -1.702028866 | biosynthesis of bacillithiol | Spx regulon, SigA regulon | yes |
| *bshB1* | -1.493706871 | biosynthesis of bacillithiol | Spx regulon, SigA regulon | yes |
| *ykuV* | -1.204354124 | protection of proteins against oxidative damage | AbrA regulon | yes |

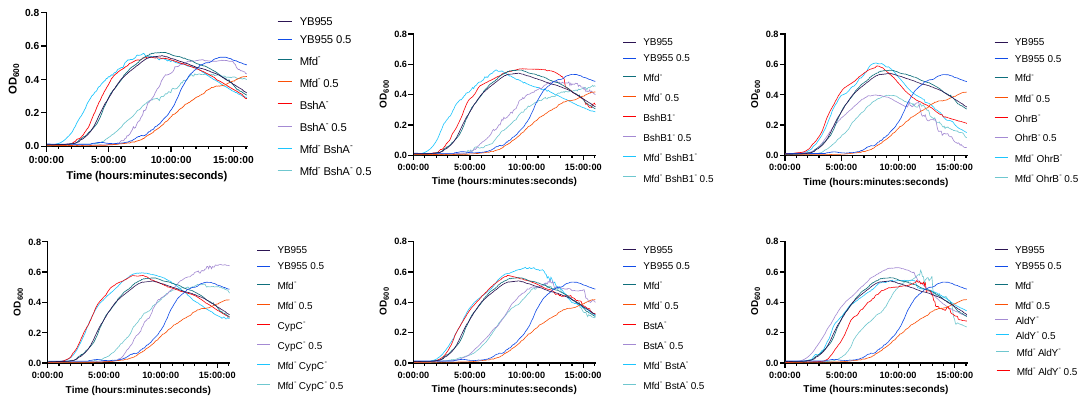

**Figure S2. Growth curves for Figure 5.** Average OD_600_ readings of strains differing in Mfd measured by a plate reader (n=9).

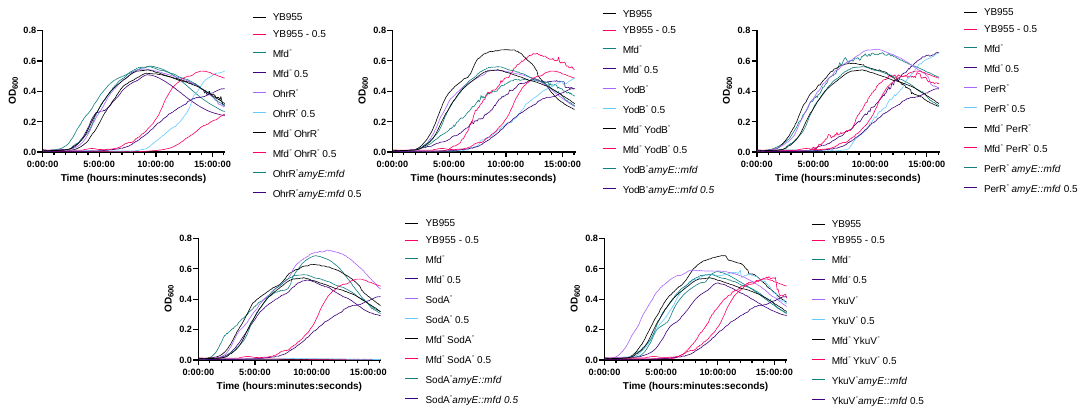

**Figure S3. Growth curves for Figure 6.** Average OD_600_ readings of strains differing in Mfd measured by a plate reader (n=9).

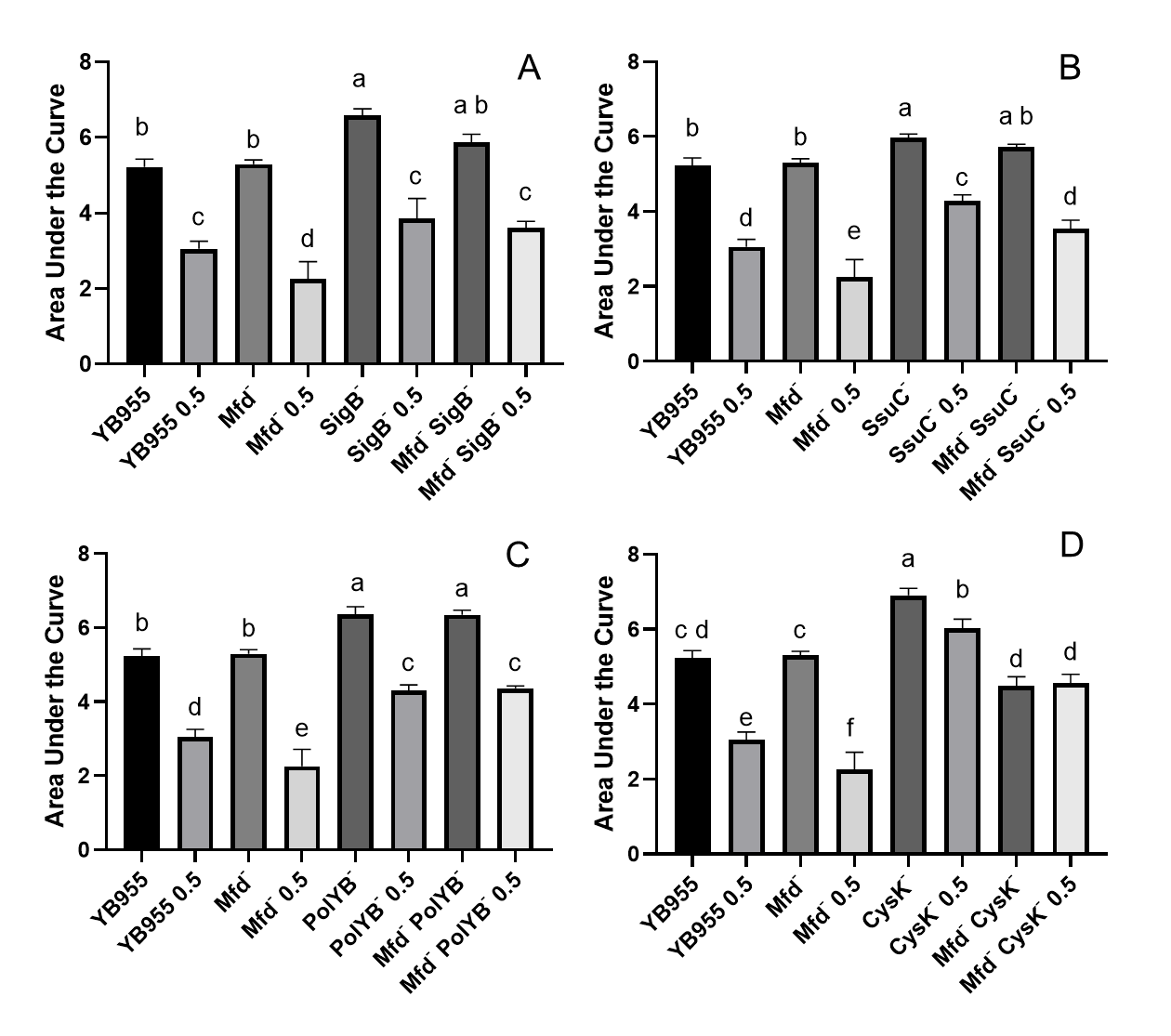

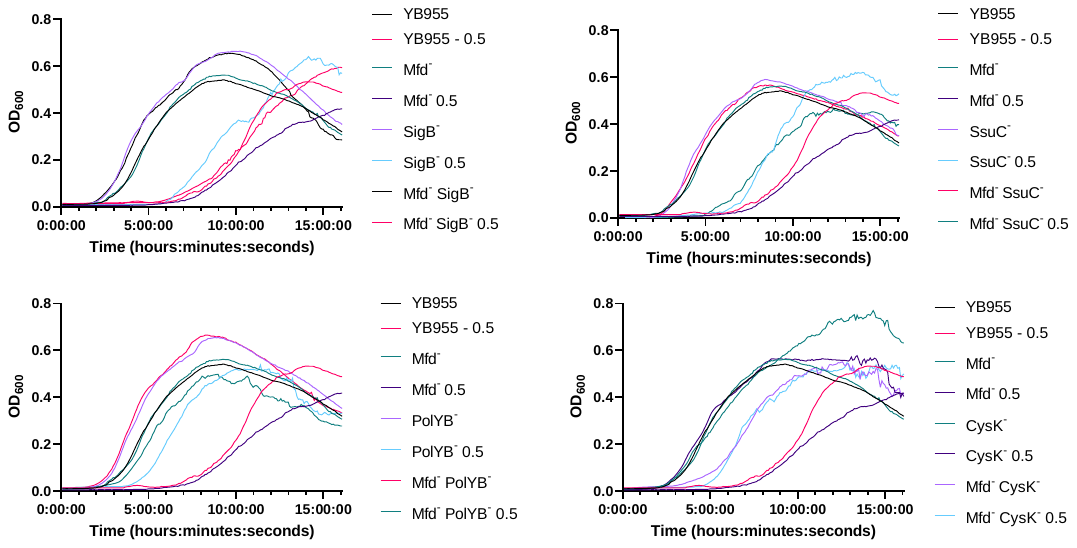

**Figure S4. Growth dynamics of single- and double-gene-deletion strains in the presence and absence of diamide. (Top)** Average area under the curve as calculated by the R package, Growthcurver, of growth curves from bottom panel. Lower case letters distinguish significant differences between means. “a”, “b”, “c” and onward are significantly different mean groups. **(Bottom)**Average OD_600_ readings of strains differing in Mfd measured by a plate reader (n=9).

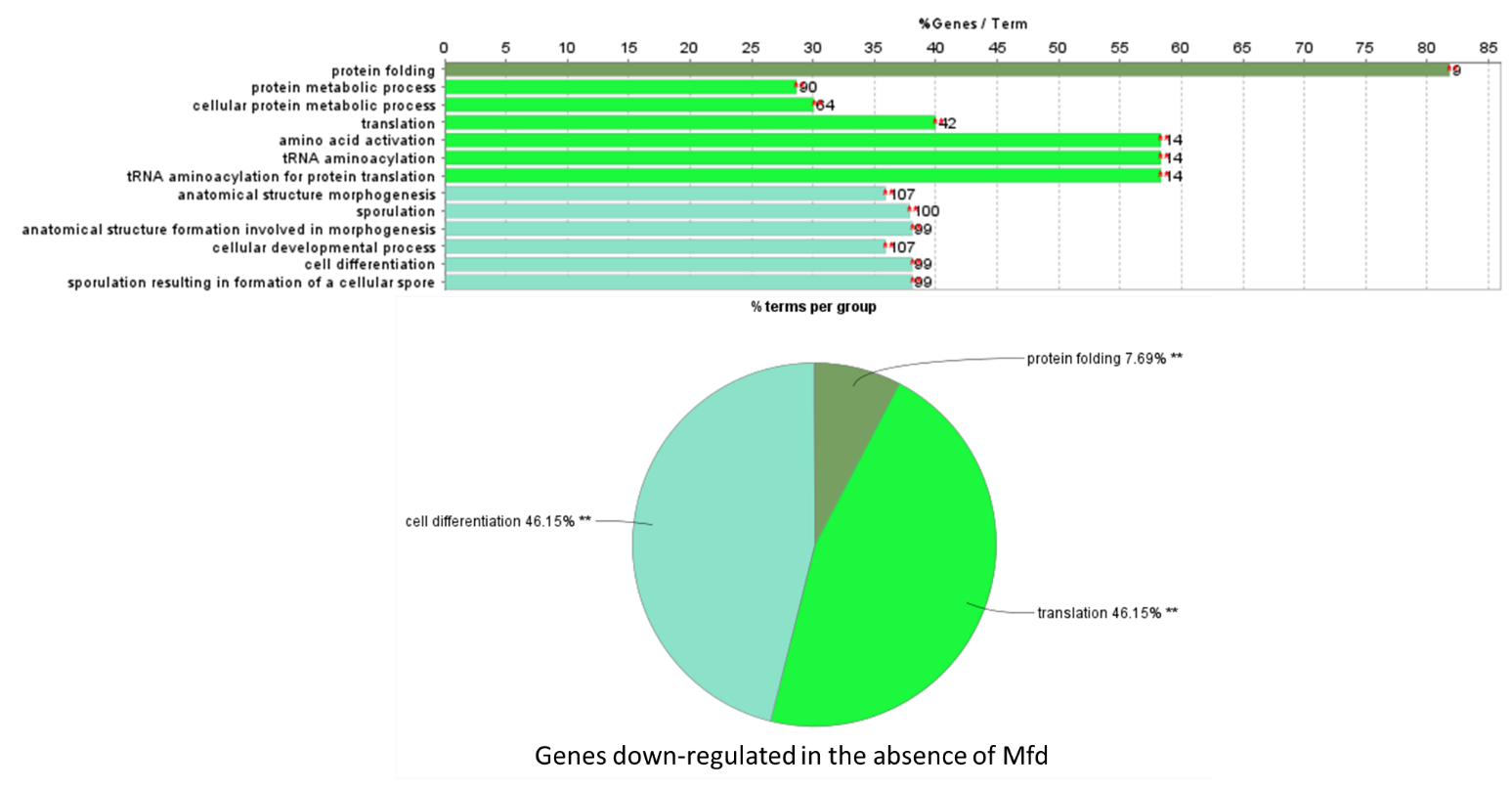

**Figure S5. Pathway enrichment analysis of down-regulated genes in the absence of Mfd in stationary-phase cells.**

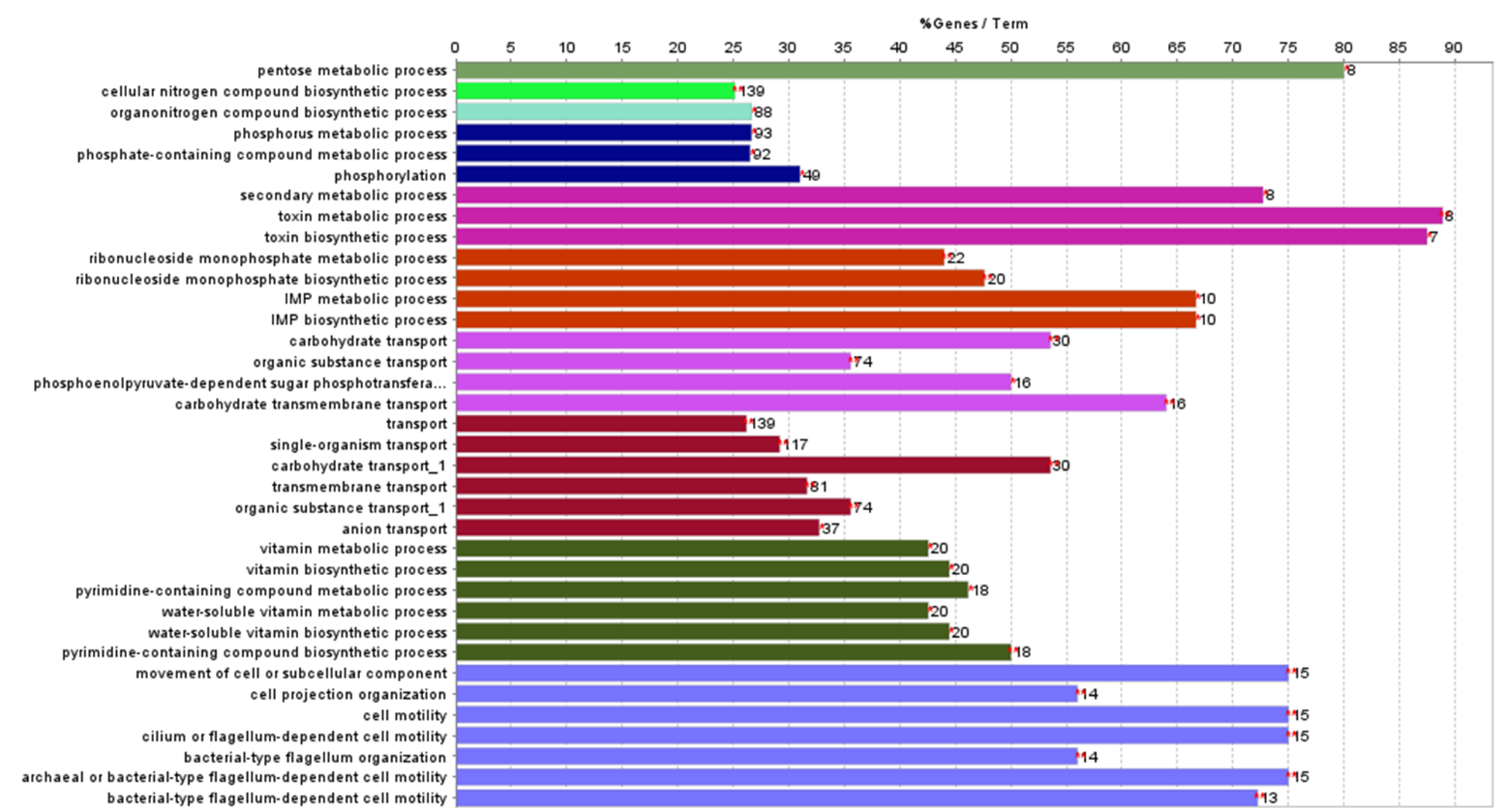

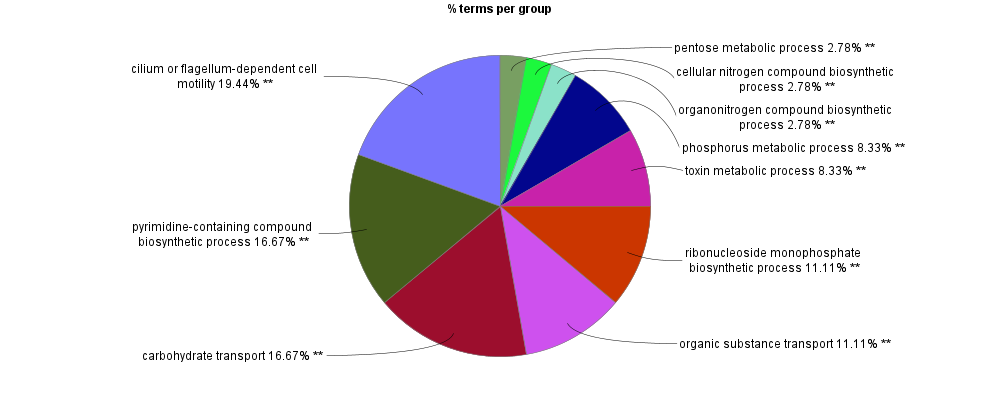

**Figure S6. Pathway enrichment analysis of genes up-regulated in the absence of Mfd in stationary-phase cells.**

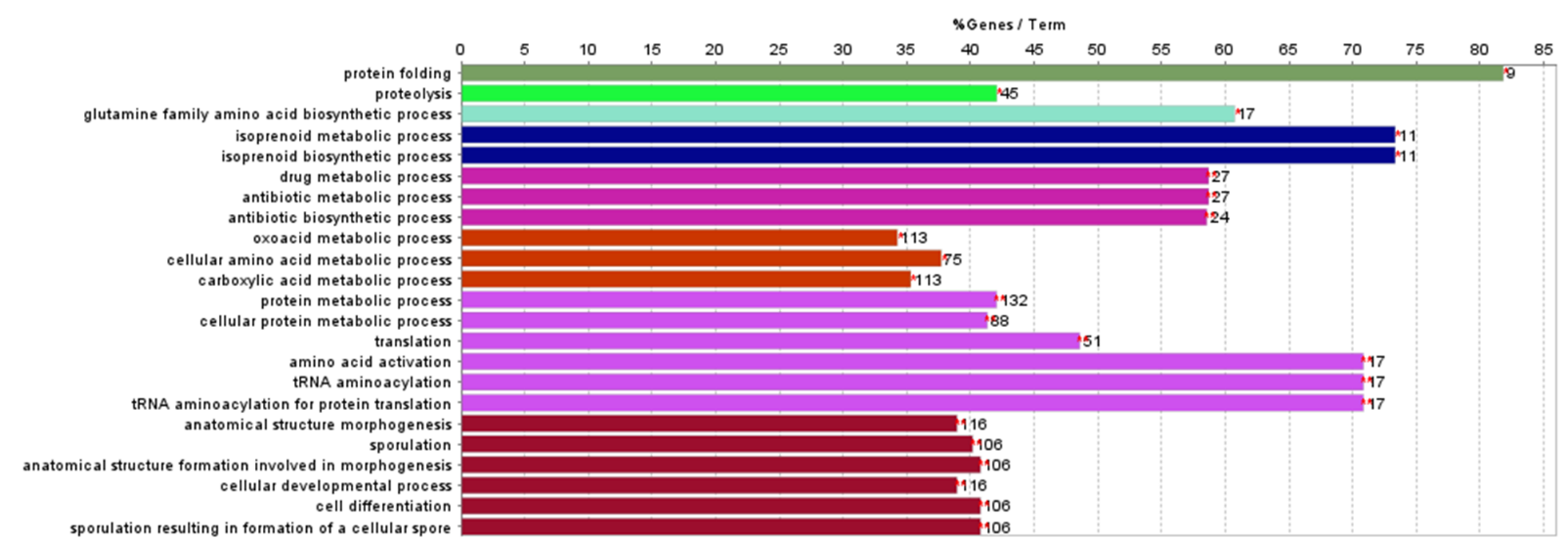

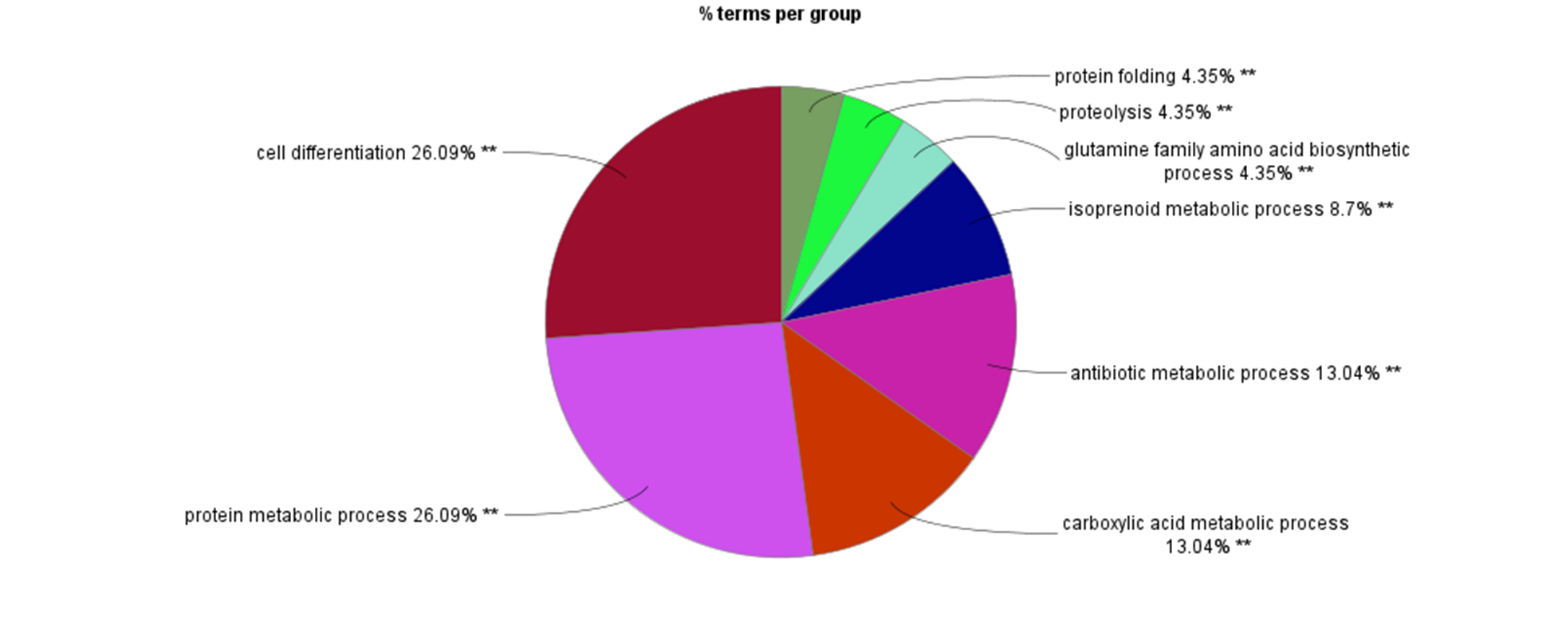

**Figure S7. Pathway enrichment analysis of genes down-regulated in the absence of Mfd and in the presence of diamide in stationary-phase cells.**

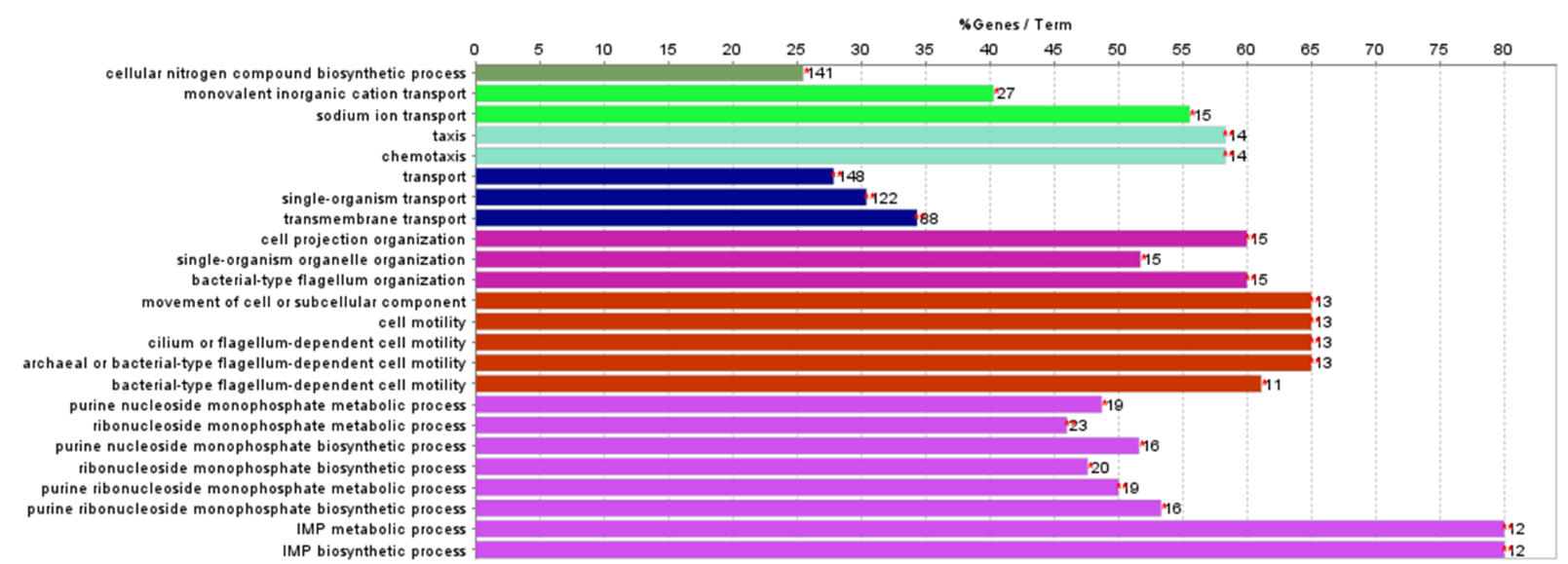

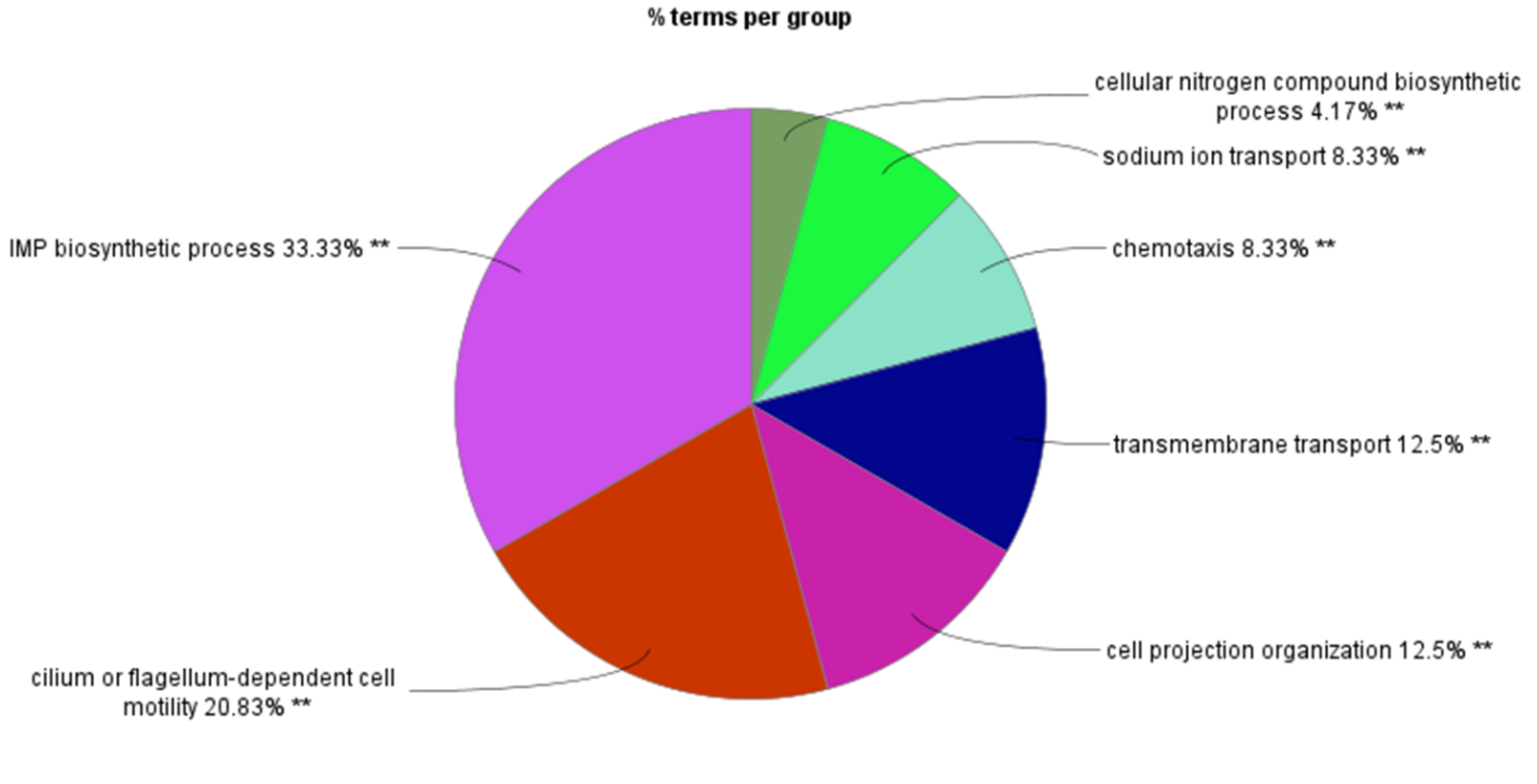

**Figure S8. Pathway enrichment analysis of genes up-regulated in the absence of Mfd and in the presence of diamide in stationary-phase cells.**
